# Cryo-electron Microscopy and Exploratory Antisense Targeting of the 28-kDa Frameshift Stimulation Element from the SARS-CoV-2 RNA Genome

**DOI:** 10.1101/2020.07.18.209270

**Authors:** Kaiming Zhang, Ivan N. Zheludev, Rachel J. Hagey, Marie Teng-Pei Wu, Raphael Haslecker, Yixuan J. Hou, Rachael Kretsch, Grigore D. Pintilie, Ramya Rangan, Wipapat Kladwang, Shanshan Li, Edward A. Pham, Claire Bernardin-Souibgui, Ralph S. Baric, Timothy P. Sheahan, Victoria D′Souza, Jeffrey S. Glenn, Wah Chiu, Rhiju Das

**Author notes:** These authors contributed equally to this work (Kaiming Zhang, Ivan N. Zheludev, Rachel J. Hagey).

## Abstract

Drug discovery campaigns against Severe Acute Respiratory Syndrome Coronavirus 2 (SARS-CoV-2) are beginning to target the viral RNA genome^1, 2^. The frameshift stimulation element (FSE) of the SARS-CoV-2 genome is required for balanced expression of essential viral proteins and is highly conserved, making it a potential candidate for antiviral targeting by small molecules and oligonucleotides^3–6^. To aid global efforts focusing on SARS-CoV-2 frameshifting, we report exploratory results from frameshifting and cellular replication experiments with locked nucleic acid (LNA) antisense oligonucleotides (ASOs), which support the FSE as a therapeutic target but highlight difficulties in achieving strong inactivation. To understand current limitations, we applied cryogenic electron microscopy (cryo-EM) and the Ribosolve^7^ pipeline to determine a three-dimensional structure of the SARS-CoV-2 FSE, validated through an RNA nanostructure tagging method. This is the smallest macromolecule (88 nt; 28 kDa) resolved by single-particle cryo-EM at subnanometer resolution to date. The tertiary structure model, defined to an estimated accuracy of 5.9 Å, presents a topologically complex fold in which the 5′ end threads through a ring formed inside a three-stem pseudoknot. Our results suggest an updated model for SARS-CoV-2 frameshifting as well as binding sites that may be targeted by next generation ASOs and small molecules.

## Introduction

Since December 2019, the SARS-CoV-2 virus, the etiological agent of Coronavirus Disease 2019 (COVID-19), has been a significant global health concern. As of early July 2020, more than 13 million people have been infected and more than 570,000 have died, as reported by the Coronavirus Resource Center of Johns Hopkins University, and this number is still growing. Effective vaccines and therapeutic drugs are needed to reduce the severity of the ongoing SARS-CoV-2 global pandemic. By early July 2020, 152 vaccines and approximately 300 therapeutic drugs are being developed worldwide for combating SARS-CoV-2 infection^8^, though most are focused on a few targets^2^. Specific drug design based on protein and RNA structures has been an effective means of drug development^9–11^. Three-dimensional (3D) structures of several key proteins of SARS-CoV-2 have been resolved^12–16^, but 3D structures of its key RNA elements remain unknown.

Like all coronaviruses, SARS-CoV-2 has a positive sense (+), single-stranded RNA (ssRNA) genome. Its first open reading frames (ORFs) 1a and 1b encode for SARS-CoV-2 non-structural proteins, including the RNA-dependent RNA polymerase, and partially overlap^17^. Optimal viral fitness is dependent on precise stoichiometric expression of ORF1a and ORF1ab throughout the viral replication cycle^17, 18^ facilitated by an essential regulatory mechanism termed −1 programmed ribosomal frameshifting (−1 PRF). The −1 PRF mechanism is stimulated by a structured RNA motif at the 3′ end of ORF1a termed the frameshift stimulation element (FSE). This element directs elongating ribosomes to stochastically shift their reading frames by one base in the 5′ direction, enabling readthrough past the ORF1a stop codon into ORF1b and maintaining appropriate levels of ORF1a to ORF1ab expression^19^.

SARS-CoV-2 has an FSE sequence identical to the original SARS-CoV-1 FSE up to a single nucleotide substitution^20^ (**Fig. 1a**, non-synonymous mutation C13533A; C63A in our numbering). As with other coronavirus FSEs, the SARS-CoV-2 FSE has a 5′ heptanucleotide ‘slippery site’ UUUAAAC followed by an RNA element hypothesized to form a three-stem pseudoknot^19, 21^. Sequence conservation, *in vitro* functional analysis, and small angle x-ray scattering (SAXS) suggest that the SARS-CoV-2 forms an FSE that folds similarly to the original SARS coronavirus FSE^3^. The SARS-CoV-2 FSE is highly conserved compared to its genomic context^22^, with few single-nucleotide polymorphisms arising in recent strains^23^, all of which sustain frameshifting in *in vitro* assays^24^. Consequently, the SARS-CoV-2 FSE appears functionally obligate for viral fitness and is predicted to have little evolutionary flexibility to evolve away from any potential therapeutics^25^. Supporting its potential as a therapeutic target, previous studies have shown that changing the FSE sequence leads to loss of viral fitness in SARS-CoV-1 and other coronaviruses^17, 18^. While secondary structures required for frameshifting have been determined by NMR and compensatory mutagenesis^25^ and probed by chemical mapping^26–28^, experimentally derived tertiary structures remain unavailable for any coronavirus FSE, and the mechanism of frameshifting remains poorly understood^25, 27, 29, 30^. Attempts to disrupt frameshifting by the SARS-CoV-1 or SARS-CoV-2 FSE with peptide nucleic acids and small molecules have provided leads for drug optimization, but these compounds have given incomplete frameshifting inactivation and not yet been tested on SARS-CoV-2 in cells^5, 6^.

**Fig. 1.**
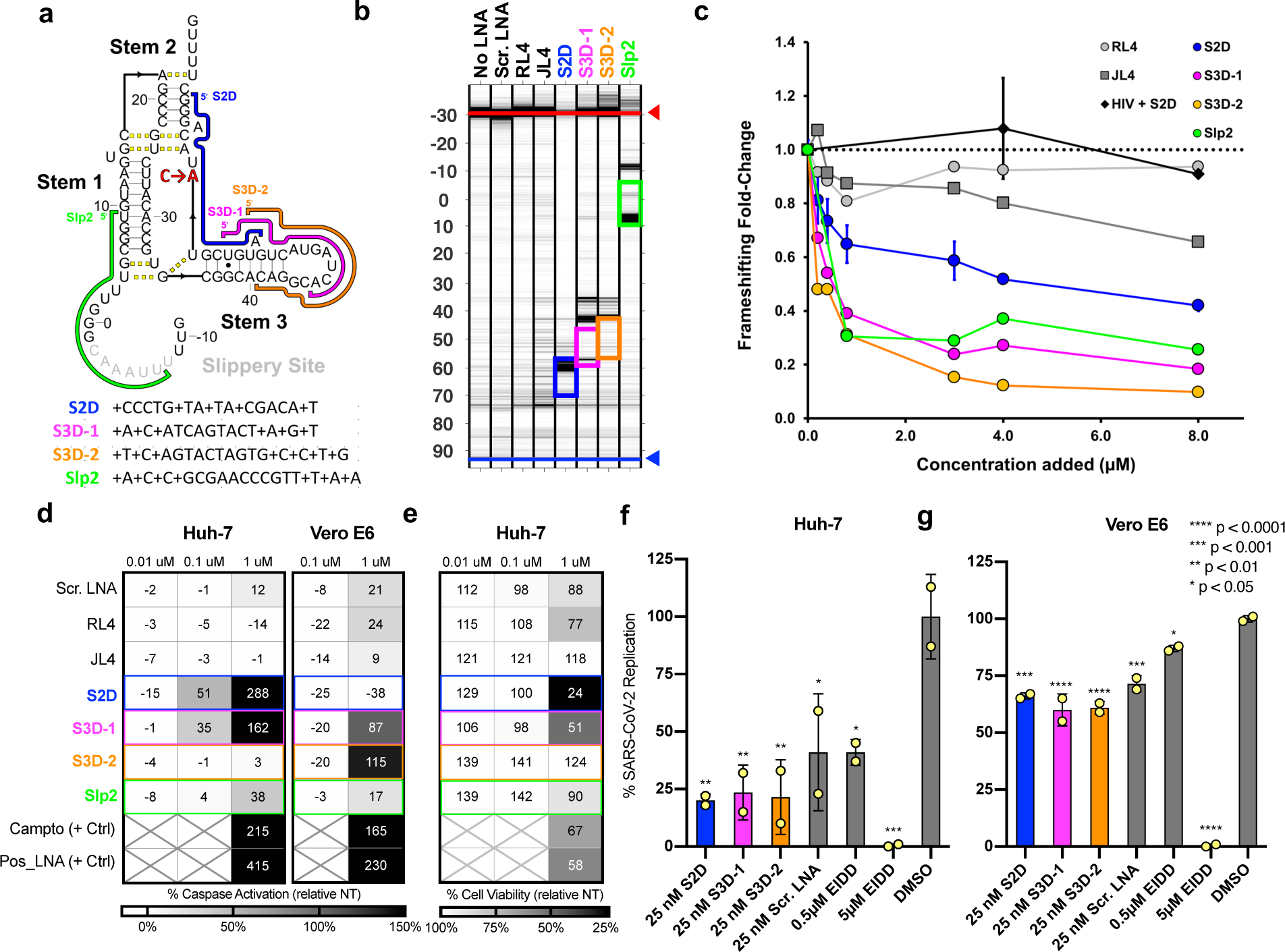
Exploratory antisense targeting of the SARS-CoV-2 FSE. **a.** Antisense oligonucleotides (ASOs) modified with LNA bases (+s) were designed against the secondary structure of SARS-CoV-2 FSE RNA. Bases with ambiguous base-pairing partners indicated with yellow dashed lines. C63A mutation between SARS-CoV-1 and SARS-CoV-2 shown in red. **b.** Capillary electrophoresis (CE) trace of reverse transcription products of SARS-CoV-2 FSE (60 nM) in the presence of LNA ASOs (1 μM). Negative controls: nontreated (no LNA), scramble (Scr.), RL4, and JL4 LNAs. Blue and red arrows indicate unextended RT primer (negligible) and fully extended cDNA, respectively. Colored boxes indicate segments at which LNA ASOs are expected to bind based on Watson-Crick complementarity. **c.** Mean *in vitro* dual luciferase frameshifting efficiencies as a function of LNA ASO concentration, shown as fold-change relative to frameshifting efficiency (26.5±0.93% for SARS-CoV-2 FSE; 5.4 ±0.05% for HIV Gag-Pol frameshift signal). **d-e**. Apoptosis induction and cytotoxicity of FSE-directed LNA ASOs in cells, evaluated through (d) caspase-3/7 activation, presented relative to non-treated, mock control in each cell-line: Huh-7 and Vero E6 cells, and (e) LNA cytotoxicity evaluated by PrestoBlue Cell Viability assay 4 days after LNA treatment in Huh-7 cells, presented relative to non-treated, mock control. Camptothecin (Campto) and caspase-inducing LNA (Pos_LNA) were used as positive controls. **f-g.** FSE-LNA inhibition of SARS-CoV-2-nLuc virus replication in (f) Huh-7 and (g) Vero E6. Cells were pretreated with 25 nM of FSE-directed LNAs or Scr. LNA 24 h prior to infection with SARS-CoV-2-nLuc reporter virus. Luciferase expression was measured 48 h post-infection. The nucleoside analog EIDD-1931^33^ was used as a positive control. Results are shown as percent luciferase expression relative to DMSO control, n = 2. *P* values were generated by GraphPad Prism software and computed as an ordinary one-way ANOVA using Dunnet′s multiple comparisons test against the DMSO control from each cell type. Error bars represent ± standard deviation.

Here, we report functional and structural results relevant for efforts to disrupt SARS-CoV-2 through FSE targeting. We report initial results of antisense oligonucleotides (ASOs) targeting SARS-CoV-2 in *in cellulo* replication assays, complemented by *in vitro* binding and frameshifting assays. These results remain exploratory due to the exigencies of university shutdowns and difficulties of measurements in biosafety level 3 (BSL3) laboratories, but we hope that release of these preliminary results will aid SARS-CoV-2 RNA targeting efforts. Then, to facilitate structure-guided drug design, we report a 3D structure of an 88-nucleotide SARS-CoV-2 FSE RNA at 6.9 Å by single-particle cryogenic electron microscopy (cryo-EM), validated by a second cryo-EM analysis involving a rationally designed RNA nanostructure. The structure shows a tertiary arrangement in which the 5′ end threads through a ring formed inside a three-stem pseudoknot. These results reveal binding pockets within the FSE structure and suggest mechanistic insights that might guide design and optimization of anti-frameshifting SARS-CoV-2 therapeutics.

## Results

### Exploratory targeting of SARS-CoV-2 FSE using locked nucleic acid antisense oligonucleotides

Previous studies have demonstrated FSE-based inhibition of virus replication with the original SARS-CoV-1 virus by perturbing the FSE primary sequence^17, 18^ and, with less efficacy, through peptide nucleic acid ASOs^5^; however, whether inhibiting the SARS-CoV-2 FSE affects viral replication has not been reported. Here, we explored the efficacy of antisense targeting strategies against SARS-CoV-2 FSE. Four ASOs modified with LNA bases were designed to hybridize complementary regions spanning the FSE, termed S2D (Stem 2 disruptor), S3D-1 (Stem 3 disruptor 1), S3D-2 (Stem 3 disruptor 2), and Slp2 (Slippery site 2) (**Fig. 1a**), along with negative control LNAs (RL4, JL4, and Scramble; Methods). We evaluated direct ASO-FSE interaction through *in vitro* binding and functional experiments. Reverse transcription interference read out by capillary electrophoresis (Methods) confirmed binding of the four FSE-hybridizing ASO at concentrations as low as 10 nM, and lack of binding of the three negative control ASOs (**Fig. 1b, Extended Data Fig. 1**). Sites of reverse transcription interference, mapped based on sequencing ladders, appeared near expected locations of ASO hybridization (**Fig. 1b**). *In vitro* dual-luciferase frameshifting assays confirmed inhibitory action of the four ASOs against the SARS-CoV-2 FSE (**Methods and Fig. 1c**), leading to up to 5-fold decreases in frameshifting efficiency, with IC_50_ values of 100-400 nM (**Extended Data Fig. 2a**). Tests with negative control LNAs RL4 and SL4 against the SARS-CoV-2 FSE and with the S2D LNA against the HIV FSE showed no significant inhibition at concentrations below 4 μM (**Fig. 1c**), confirming specificity of ASO-RNA interaction.

**Fig. 2.**
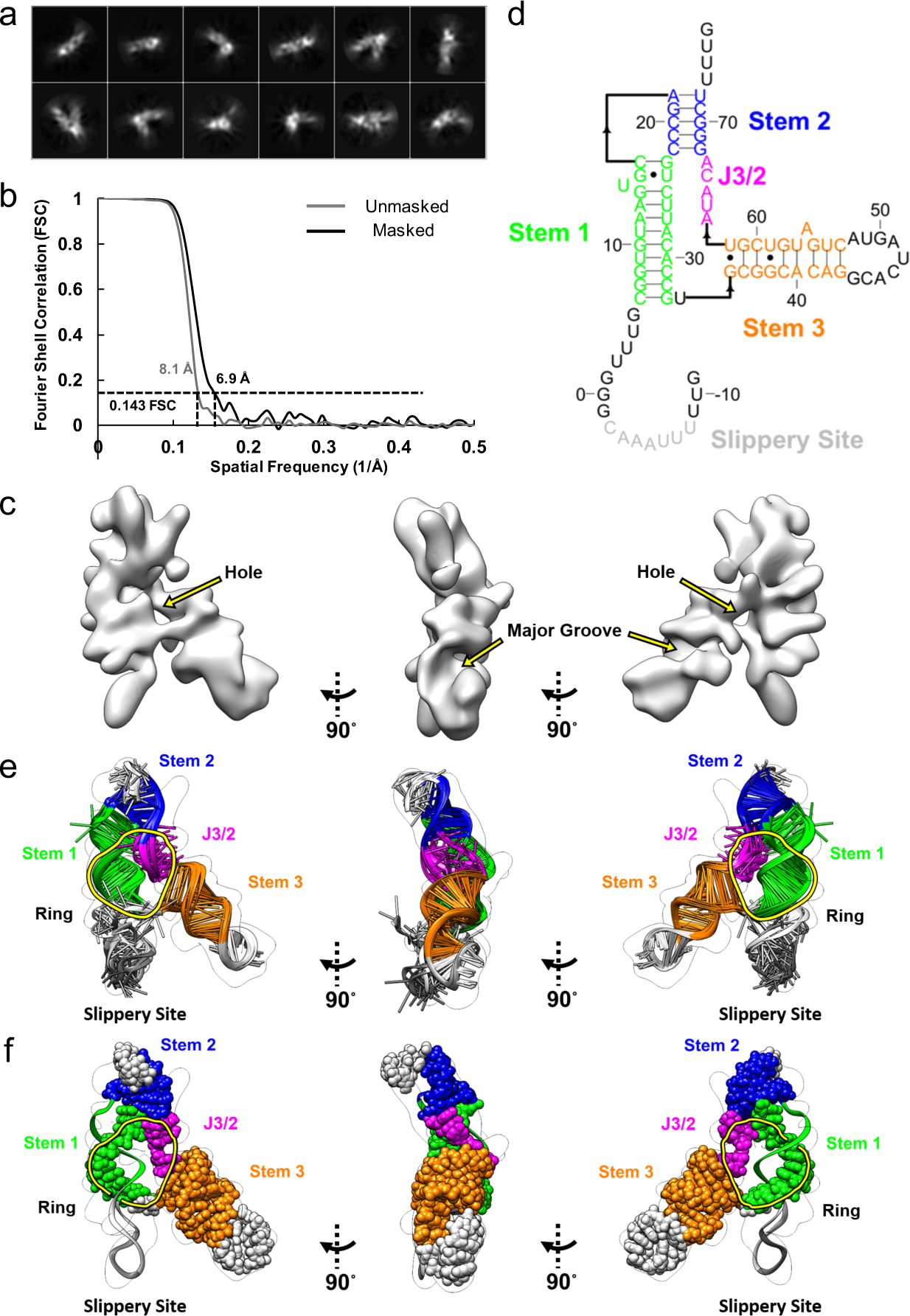
Single-particle cryo-EM analysis and model building of the FSE. **a.** Reference-free 2D class averages. **b.** Gold standard FSC plots calculated in Relion. **c.** Reconstructed cryo-EM map in three different views. **d.** Secondary structure of the FSE as determined by 1D SHAPE chemical mapping. **e.** Top 10 tertiary structures of the FSE as determined by autoDRRAFTER using the secondary structure from (c), where equivalent structural elements are indicated. These 10 top scoring models had a mean pairwise root mean squared deviation of 5.68 Å, resulting in an estimated accuracy of 5.9 Å based on the previously determined linear relationship7. **f.** A ring formed by the second strand of Stem 1 with Stems 2 and 3 and J3/2, into which the first strand of Stem 1 threads to form a topologically constrained tertiary fold.

In parallel to these cell-free assays, we tested the antiviral potential of FSE-directed LNA ASOs through cellular assays in the human liver cell-line Huh-7 and the African green monkey kidney cell-line Vero E6. To evaluate cytotoxic effects of the ASOs, we screened each ASO for both cellular caspase-3/7 activation, as a measure of apoptotic induction^24^, and cell viability over 4 days. Nearly all ASOs were insignificant activators of caspase and were not cytotoxic at concentrations of 100 nM or below (**Fig. 1d and Fig. 1e**). To test antiviral potency, we measured the effects of the ASOs on a replicating SARS-CoV-2 luciferase reporter virus (SARS-Cov-2-nLuc) under BSL3 conditions. Cells were treated with S2D, S3D-1, and S3D-2 LNAs at a concentration of 25 nM, well below their cytotoxic concentrations and within the therapeutic dose range typical of potent LNA inhibitors in cell culture^31, 32^ (**Fig. 1e**). Compared to the DMSO control, S2D, S3D-1, and S3D-2 displayed statistically significant virus inhibition by approximately 4-fold in Huh-7 cells (p < 0.01; **Fig. 1f; Extended Data Fig. 2b**). These ASO effects at 25 nM were also stronger than those of 25 nM scrambled LNA negative control and 0.5 μM positive control compound, EIDD-1931 (NHC, β-d-N4-hydroxycytidine^33^), though current variabilities in transfection leave these differences statistically insignificant. Similar results, with less variability between replicates but overall weaker reductions, were observed in Vero E6 African green monkey kidney cells (**Fig. 1g; Extended Data Fig. 2c**).

These exploratory experiments motivate moving forward with viral replication measurements involving variation of ASO concentrations, mutagenesis to test specificity, and tests with Slp2 and other ASOs, which are being scheduled at BSL3 facilities. In particular, we cannot presently rule out non-specific inhibition by LNA ASO transfection. Nevertheless, taken together, our binding, frameshifting, and viral replication assays are consistent with FSE as a possible therapeutic target and provide candidate ASO molecules and assays for follow-on characterization and optimization. In our cell-free experiments as well as other studies on ASOs and small molecules targeting SARS-CoV-1 and SARS-CoV-2 frameshifting^5, 34^, saturating concentrations of candidate anti-frameshifting inhibitors have shown only partial inactivation (2- to 5-fold), whereas compounds targeting other viral mechanisms reduce viral replication by 100-fold or more at high concentrations (5 μM EIDD-1931, **Fig. 1f-g**). We hypothesized that the partial inactivation and modest affinity of these targeting efforts so far might be better understood and eventually overcome if we could achieve a better picture of the tertiary structure of the SARS-CoV-2 FSE, discussed next.

### Cryo-EM resolves the 88-nt FSE

Cryo-EM structure determination of RNA-only systems or macromolecules with molecular weights under 50 kDa has been accomplished only recently^7, 35–38^, and it was unclear whether cryo-EM might be applicable to an RNA as small as the SARS-CoV-2 FSE (28 kDa, 88 nucleotides; **Fig. 1a**). Initially, we collected a dataset comprising ∼13,000 micrographs. However, the low concentration of the sample on the grids and the high background of molecular species in the raw images made it difficult to perform image processing (**Extended Data Fig. 3**). We manually selected ∼3,400 images out of the whole dataset with the defocus higher than −3 μm with the aim of achieving an initial 3D reconstruction from ∼27,000 manually picked particles (**Extended Data Fig. 3**). Subsequently, we optimized the sample preparation and RNA folding protocol to improve the stability and the concentration of stably folded particles on the electron microscope grids (**see Methods**). A dataset with ∼10,000 micrographs was collected using the optimized sample (**Extended Data Table 1**). Benefiting from the lower background, about one million particles were selected with the Neural Network particle autopicking option in EMAN2^39^ and were subject to 2D and 3D image classification in Relion^40^ (**Extended Data Figs. 3 and 4**). We selected one class of images yielding a reconstruction with well-connected density for further refinement. The reference-free 2D classification of this particle set and its refined 3D map display a shape similar to the Greek character “λ” (**Fig. 2 and Extended Data Fig. 3**) and the final resolution is 6.9-Å (**Fig. 2b**). The B-factor^41, 42^, calculated based on a fitted relationship of particle number and map resolution, was estimated to be ∼726 Å^2^ (**Extended Data Fig. 5**). Such a large B-factor could be attributable to the flexibility of the molecule and/or the limit of particle orientation estimation accuracy due to its small size. Nevertheless, at the current 6.9-Å resolution, the map still permitted identification of helical groove features that are hallmarks of RNA (**Fig. 2c**). The map also showed non-helical crevices and holes suggestive of potential small molecule binding pockets (**Fig. 2c**), as previously observed and validated in cryo-EM maps of SAM-IV riboswitch aptamers with and without small molecule ligands^7, 37^.

**Fig. 3.**
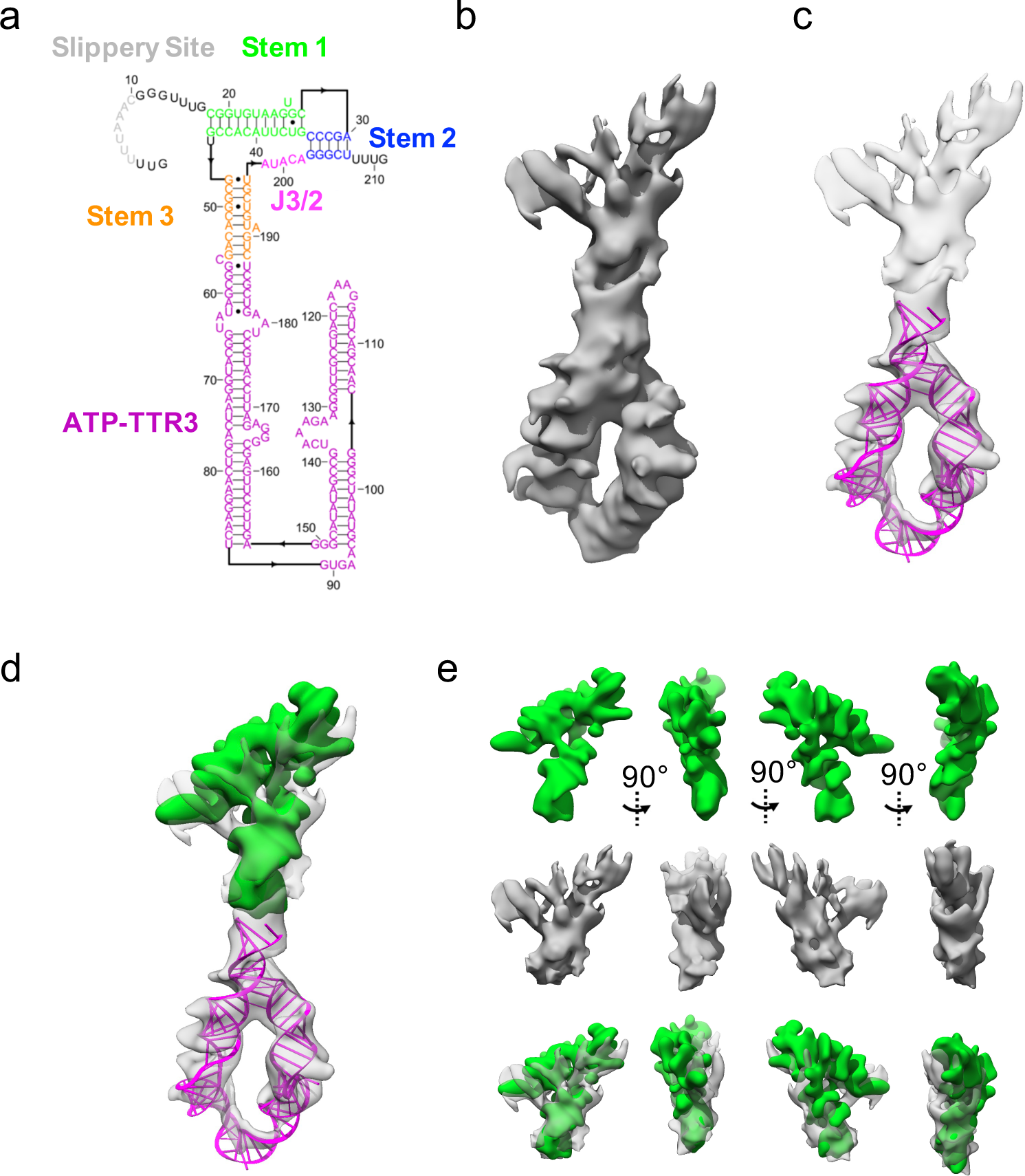
Single-particle cryo-EM analysis of nanostructure-tagged FSE. **a.** Secondary structure of the FSE-ATP-TTR3 as determined by SHAPE chemical mapping, with key structural elements indicated in different colors. **b.** Cryo-EM map of the FSE-ATP-TTR3 at a low contour level, oriented as in (a)**. c.** ATP-TTR 3 model (PDB code: 6WLK) docked into the FSE-ATP-TTR3 map. **d.** FSE map from Fig. 2., green, docked into the extra density of the FSE-ATP-TTR3 map. **e.** Comparison between the FSE map and the extra density of the FSE-ATP-TTR3 map.

**Fig. 4.**
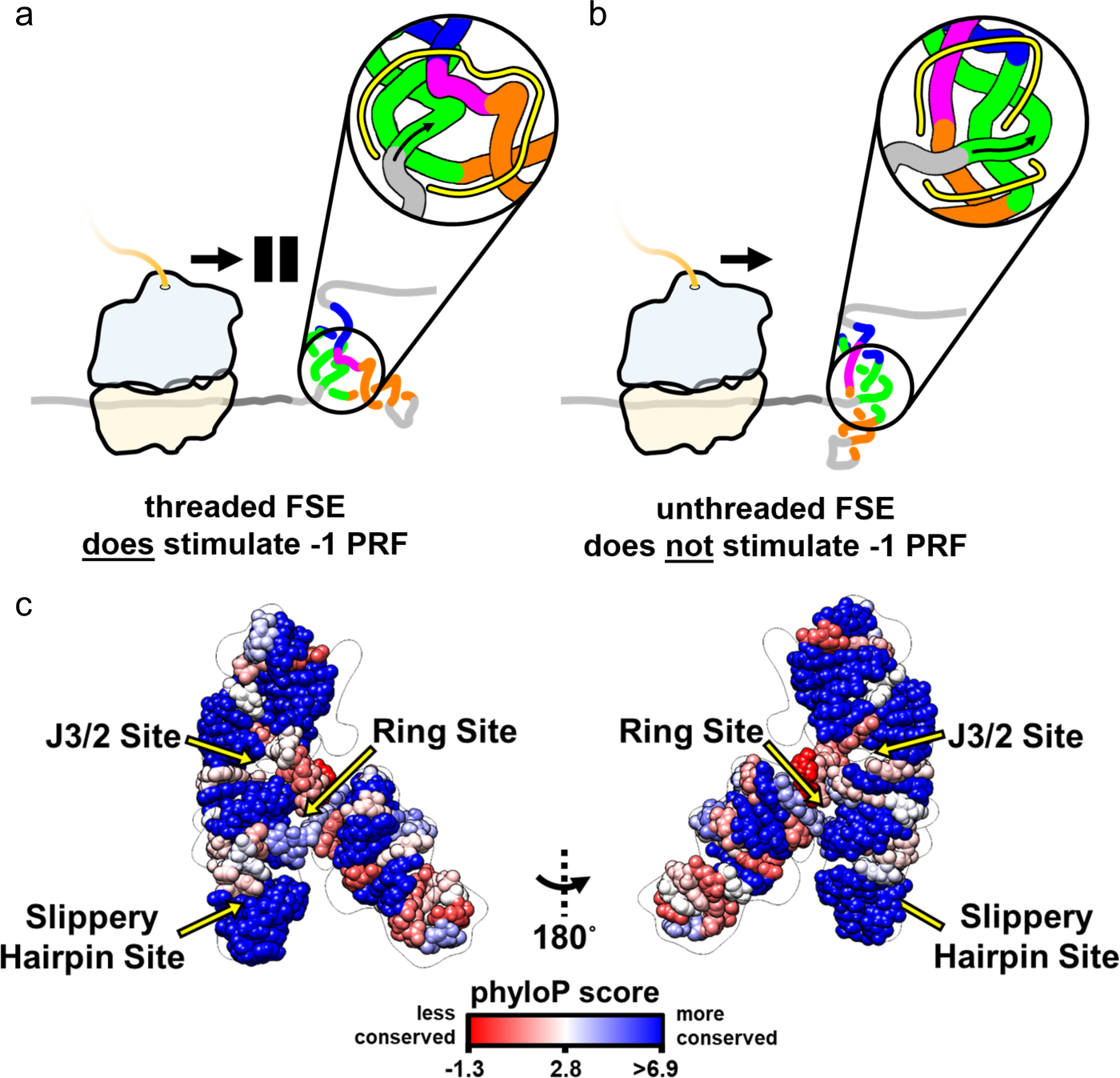
Structural complexity in SARS-CoV-2 −1 programmed ribosomal frameshifting. After the ribosome unfolds upstream genomic structures, the FSE refolds, resulting in a 3D fold in which the 5′-end is either (**a**) threaded through a ring in the FSE pseudoknot (inset, hole indicated in yellow), as captured in our cryo-EM structure, or (**b**) left unthreaded. The model in (**b**) reflects an alternative fold with the same pseudoknotted secondary structure but distinct tertiary arrangement, as captured by *de novo* modeling. The torsional restraint^61^ model suggests that the cryo-EM-resolved conformation in (**a**) promotes pausing at the slippery site and ribosomal frameshifting and should be targeted for destabilization relative to alternative folds (**b**) in antiviral efforts seeking to disrupt frameshifting, or stabilization in efforts that seek to increase frameshifting. **c.** Three potential small molecule binding sites suggested by the cryo-EM-resolved structure involve nucleotides that are highly conserved relative to the rest of the genome. Model coloring is by the phyloP score derived from 119 diverse coronaviruses^22, 77^, with blue vs. red showing higher vs lower conservation, respectively, compared to the mean phyloP score of the entire SARS-CoV-2 genome.

### Ribosolve modeling of FSE coordinates

To better understand the FSE tertiary structure, we sought to model RNA coordinates into the 6.9-Å map using Ribosolve, a hybrid pipeline recently developed for automatically modeling RNA 3D structures based on secondary structure information from mutate-and-map guided by next-generation sequencing (M2-seq), cryo-EM maps, and computer modeling with autoDRRAFTER^7^. M2-seq secondary structure analysis recovered the three-stem pseudoknot in Fig. 1a that has been validated by NMR and compensatory mutagenesis for the SARS-CoV-1 FSE^21^. The same three-stem secondary structure was also observed in a separate SHAPE-directed modeling study as well as an independent DMS-MaPseq study^27^, with minor variations in edge base pairs (**Extended Data Fig. 6**). Using the consensus secondary structure of **Fig. 2d** and the cryo-EM map, autoDRRAFTER modeling produced an ensemble of ten structures with a mean pairwise root mean squared deviation (RMSD) of 5.68-Å (**Fig. 2e, Extended Data Fig. 7 and Extended Data Movie 1**), corresponding to a 5.9-Å estimated accuracy, based on a linear relationship of precision and accuracy discovered previously^7^, with the largest uncertainty in the 5′ end and at Stem 2 (estimated accuracy of >10 Å; **Extended Data Fig. 8**). The quality of both model ensembles was also validated using MolProbity^43^ (**Extended Data Table 1**). At 5.9-Å estimated accuracy, individual atomic positions and non-canonical base pairs cannot be confidently assigned. Nevertheless, the tertiary arrangement of the helical segments and non-helical linkers of the SARS-CoV-2 FSE can be traced (**Fig. 2**), is consistent in different members of the model ensemble, and is further supported by alternative autoDRRAFTER runs making different assumptions about secondary structure and initial 3D helix placements (**Extended Data Fig. 7**).

### Overall architecture of the SARS-CoV-2 FSE

The architecture of the FSE involves several interlocking elements (**Fig. 2e-f and Extended Data Movie 1**). Starting from the 5′ end and proceeding to the 3′ end, the molecule begins with a 5′ region that includes the heptanucleotide slippery site. The Ribosolve modeling folds this region into a loose hairpin-like shape closed by G•U wobble pairs. This 5′ end is followed by the first strand of Stem 1, a long helix in all coronavirus FSE’s. The loop of Stem 1 is also the first strand of the Stem 2 pseudoknot, which forms a 5-bp helix that hybridizes with its complement at the 3′ end of the FSE. The RNA strand continues from this region to complete the second strand of Stem 1 and doubles back to form a hairpin, Stem 3. After an unpaired segment J3/2, the RNA completes Stem 2 to close the Stem 1-Stem 2 pseudoknot. Lastly, unstructured terminal nucleotides form a 3′ tail.

Despite the absence of base pairings or direct stacking between Stem 3 and the Stem 1-Stem 2 pseudoknot, Stem 3 exhibits a distinct tertiary conformation in relation to the pseudoknot which, along with the conformationally heterogeneous hairpin at the 5′ end, result in the legs of the “λ”-shaped map (**Fig. 2e-f, Extended Data Fig. 6 and Extended Data Movie 1**). Overall, Stems 1, 2, and 3 form a circular ring with a visually apparent hole (**Fig. 2e-f**). The 5′ end of the FSE is connected to the pseudoknot by a linker that passes through this ring. This complex topology was predicted as a possible FSE fold in two recent, independent 3D computer modeling studies that include an extended 5′ end^44, 45^. We emphasize that while the fold of the 5′ end appears poorly resolved and thus may have multiple conformations, the connection point of the 5′ end to the rest of the structure and its threading through the Stem 1-Stem 2-Stem 3 ring is a consistent feature in all models in the autoDRRAFTER ensemble (**Extended Data Movie 1**) as well as in alternative modeling runs based on alternative FSE secondary structures and different autoDRRAFTER modeling assumptions (**Extended Data Fig. 7**). In terms of structural requirements, the 5′ end ring-threading requires formation of a ∼10 base pair helical turn in Stem 1. With fewer base pairs, the 5′ strand of Stem 1 cannot turn fully inside and through the ring. Supporting the general relevance of 5′-end ring-threading, the length of Stem 1 is ∼10 base pairs or larger in all proposed coronavirus FSE elements^22^, and *de novo* modeling of FSEs from bovine coronavirus, murine hepatitis virus, human coronaviruses OC43 and HKU1, SARS-CoV-1, and MERS all give 5′ ring-threaded models (**Extended Data Figure 9**).

### Validation of the FSE model through RNA nanostructure tagging

The SARS-CoV-2 FSE RNA represents an extreme case for cryo-EM. It is the smallest macromolecule (28 kDa) so far resolved by cryo-EM single particle analysis at subnanometer resolution. We therefore sought further independent validation of the map and the Ribosolve model, particularly to test our inference that Stem 3 comprises a ′leg′ of the λ shape, perpendicular to the Stem 1-Stem 2 pseudoknot^44, 45^. We rationally designed a variant of the SARS-CoV-2 FSE termed FSE-ATP-TTR3, which contains an insertion of a clothespin-like nanostructure whose visualization would test the assignment and orientation of Stem 3 in the FSE map^46^ (**Fig. 3a**). For the tag, we chose the rationally designed ATP-TTR 3 RNA^47^ based on its known amenability to cryo-EM imaging^7^ and on modeling suggesting that its insertion into the FSE would not perturb either RNAs’ secondary structure, a prediction verified through chemical mapping (**Extended Data Fig. 10**).

A dataset containing ∼12,000 micrographs of FSE-ATP-TTR3 was collected and resulted in a 6.4-Å map of FSE-ATP-TTR3 exhibiting the ATP-TTR 3 clothespin-like shape. Additional density at the end of the clothespin is visible, with a λ-like shape (**Fig. 3b-c, Extended Data Fig. 11 and Extended Data Movie 2**). The FSE-only map fits into this additional density (**Fig. 3d, Extended Data Movie 2**). To evaluate the relative orientation of FSE and the ATP-TTR tag, we adopted an unbiased method to align the two maps. The orientation of the FSE map was determined by conducting a rigid body exhaustive search to maximize the correlation between the FSE density and difference mapping between FSE-ATP-TTR3 and ATP-TTR3 densities independently using Segger^48^ and Situs^49^. The highest cross-correlation was found at the expected site, confirming the orientation and location of Stem 3 within the FSE map (orange helix, **Fig. 3e; Extended Data Fig. 12 and Extended Data Movie 2**) and prospectively corroborating the prediction from autoDRRAFTER analysis of the FSE alone.

## Discussion

The urgency of the COVID-19 pandemic, recent advances in targeting RNA 3D structures with ASOs and small molecules, and the identification of the FSE as a potentially well-defined RNA 3D structure in the SARS-CoV-2 genome has generated significant interest in understanding and targeting SARS-CoV-2 ribosomal frameshifting^3, 23, 27, 28, 44, 45, 50, 51^. Here, we confirmed the ability of ASOs to invade the SARS-CoV-2 FSE structure and to reduce frameshifting efficiencies in cell-free assays, and we explored the potential for these ASOs to reduce viral replication in cells at sub-micromolar concentrations. Incomplete inactivation of coronavirus frameshifting and replication in these and prior studies motivated structural analysis to better understand the FSE. Despite prior expectations of structural heterogeneity and propensity for multimerization^4, 45, 52^, and despite having a size slightly under that of the previous smallest macromolecule imaged by cryo-EM (28 vs. 30 kDa^35^), the SARS-CoV-2 FSE gave a 6.9-Å resolution monomer map with recognizably helix-like elements arranged into a λ-like tertiary arrangement. Automated secondary structure determination and 3D coordinate building through the recently developed Ribosolve pipeline led to a model with an intricate fold: the Stem1-Stem2 pseudoknot interconnected by a J3/2 linker and Stem 3 to form a closed ring. Remarkably, the linker connecting the 5′ end of the FSE to Stem 1 threads through this ring. This overall 3D arrangement was tested through a second cryo-EM analysis of an FSE molecule extended by a rationally designed clothespin-shaped nanostructure at Stem 3.

We note two caveats regarding our cryo-EM analysis. First, the 6.9 Å resolution of the cryo-EM map is not sufficient to directly resolve base interactions, much less atom positions. It is likely that hydrogen bonds and base pairs beyond our modeled secondary structure stabilize the FSE 3D structure, and higher resolution structures will be needed to resolve those details. Second, cryo-EM data processing can filter out alternative, less well-defined conformations while focusing on particle images that contribute to the most well-defined structures. *De novo* computer modeling, in addition to predicting the ring-threaded structure we observe, has suggested alternative conformations in which the 5′ end is not threaded through the ring (**Extended Data Figure 9**). Such alternative states of FSE may exist in our preparations but escape computational detection due to the extra conformational heterogeneity at the 5′ ends. Within our nanostructure-tagged map (**Fig. 3b**), alternative states may contribute to the lower resolution of the FSE segment compared to the nanostructure segment.

Our cryo-EM-guided data and modeling, along with recent results from several groups, suggest explanations for why FSE targeting efforts by us and others have so far yielded only modest inhibition of frameshifting and viral replication^4–6, 34^. There is accumulating evidence that, in the full SARS-CoV-2 genome context, the FSE forms alternative structures involving genomic segments upstream of the element (**Extended Data Fig. 13** and refs^27,44, 45, 50^). As the human ribosome approaches the FSE, it must unfold these upstream structures through its helicase activity, and the FSE will refold. The position of the 5’ end of the FSE will be influenced by structures that existed prior to ribosome-induced refolding and could end up either threaded through the final FSE Stem1-Stem2-Stem3 ring, as captured by our cryo-EM analysis, or remain unthreaded (**Fig. 4**). The 5’-end threaded structure (**Fig. 4a**) appears poised to lead to ribosomal pausing and −1 PRF through a torsional restraint mechanism^21^, perhaps working in concert with specific interactions of the pseudoknot with the ribosome^53^. Other structures (**Fig. 4b** shows one model) might be resolved by the ribosome without frameshifting. Further work, potentially involving design of topologically trapped FSE variants, single molecule FRET, and time-resolved cryo-EM of the ribosome^54–57^, will be necessary to test and refine this model.

The structural complexity of FSE 3D folding has implications for therapeutic targeting efforts. First, for ASO targeting, the intricately threaded FSE tertiary structure (**Fig. 4c**) may hinder strand invasion, even into regions typically considered unpaired, thus explaining the requirement we and others have observed for near-micromolar ASO concentrations to achieve inhibition of frameshifting. Second, for either ASO or small molecule inhibitors, if the molecule does not significantly alter the partitioning in how the FSE 5’-end leads into the pseudoknot (**Fig. 4a vs. 4b**), it would not alter frameshifting efficiencies to extremely low levels, as is expected to be needed to disrupt viral replication^51^. If this explanation is correct, design of anti-frameshifting therapeutics may need to focus on binding to alternative structures that are distinct from our cryo-EM-guided model. Alternatively, SARS-CoV-2 replication might be disrupted by maximizing rather than inhibiting frameshifting, which would still perturb the stoichiometry of viral proteins^17, 18^. In this largely unexplored strategy, pro-frameshifting drugs could be designed to stabilize binding pockets that consistently appear in all members of the cryo-EM-guided model ensemble (**Extended Data Figure 14**). Three such sites, which we term the ‘ring site’, the ‘J3/2 site’, and the ‘slippery hairpin site’, involve nucleotides that are highly conserved across a diverse range of coronaviruses (blue in **Fig. 4c**) and hence might be targeted with a reduced chance of viral escape.

We have provided the first 3D structural data of a functionally obligate segment of the SARS-CoV-2 genome and have described implications for mechanism and targeting of −1 programmed ribosomal frameshifting, an intricate and critical genomic process. Dozens of other segments of the 30 kb RNA genome are highly conserved and have been predicted to be structured^27, 44, 45, 50, 58, 59^, many in the size range explored here. Applying antisense targeting and cryo-EM to these segments may yield additional information that sheds light on poorly understood SARS-CoV-2 RNA biology and, hopefully, accelerates design of genome-disrupting therapeutic agents.

## Materials and methods

### Locked Nucleic Acid (LNA) design and preparation

LNAs are oligonucleotides with a 2′ sugar modification that locks the ribose in a C3′-*endo* conformation by a 2′-O, 4′-C methylene bridge, which in turn locks the LNA into the A-form helical conformation^60^. Oligonucleotides containing LNAs were custom synthesized by Integrated DNA Technologies, purified by high performance liquid chromatography, and resuspended in ddH_2_O. (+) denotes LNA base. All others represent (non-locked) DNA nucleotides. Select LNA ASOs made with phosphorothioate (PS) internucleotide linkages as indicated below.

**S2D (Stem 2 Disruptor): 5′** +CCCTG+TA+TA+CGACA+T **3′**

**S3D-1 (Stem 3 Disruptor-1): 5′** +A+C+ATCAGTACT+A+G+T **3′**, PS linkages

**S3D-2 (Stem 3 Disruptor-2): 5′** +T+C+AGTACTAGTG+C+C+T+G **3′**, PS linkages

**Slp2 (Slippery site 2): 5′** +A+C+C+GCGAACCCGTT+T+A+A **3′**, PS linkages

*Negative control LNA ASOs:*

**Scramble LNA (Scr. LNA): 5′** C+ATAC+GTC+TAT+AC+GCT **3′,** PS linkages

**JL4: 5′** +CCCTG+TAAA+ACGGG+C **3′**

**RL4: 5′** +CGGGC+TAAA+AGCCC+T **3′**

*ASO control for caspase assay:*

**Pos_LNA: 5′** +A+G+C+CAGACAG+C+G+A **3′**, PS linkages

The JL4 and RL4 ASOs, inspired by ref.^61^, contained short regions of hybridization to Stem 1 and Stem 2, but were not expected to be able to displace intramolecular pairings in those stems.

### Reverse transcription interference assays

To test binding of each LNA to the SARS-CoV-2 FSE, we initially planned to carry out SHAPE and DMS chemical mapping but discovered in preliminary experiments that the bound S2D LNA interfered with reverse transcription. We therefore used this interference itself to test for binding. Binding assays were performed in 20 μL reactions containing either 1.2 pmol (0.06 μM RNA in 20 μL) or 0.2 pmol (0.01 μM in 20 μL). Starting in 4 μL of ddH_2_O, the RNA was incubated at 90°C for 3 mins, then cooled down to room temperature. Next, 4 μL of 5⨉ First strand buffer (1⨉ First strand buffer: 50 mM Tris-HCl pH 8.3, 75 mM KCl, 3 mM MgCl2) was added and incubated at 50°C for 30 mins, cooled down to room temperature prior to adding of 2.0 μL of either 0.0, 0.1, 1.0 and 10.0 μM of LNA, and further incubated at 37°C for 15 mins. Next, 5 μL of Oligo-dT-bead mixture, containing 1.5 μL of Oligo dT beads (poly(A) purist, Ambion), 0.25 μL of 0.25 μM of FAM-labelled Tail2 primer for 1.2 pmol RNA or 0.042 μM of FAM-labelled Tail2 primer for 0.2 pmol RNA, and 3.25 μl ddH_2_O, was added. Lastly, a premixed volume containing 1.0 μL of 100 mM DTT, 0.5 μL of SuperScript III reverse transcriptase (Thermo Fisher Scientific), 1.6 μL of 10 mM dNTPs and 1.9 μL of ddH_2_O was added to each reaction and reverse transcription was carried out at 48°C for 40 mins. The resulting products were then treated equivalently to the standard 1D SHAPE/DMS chemical mapping protocol (described below).

### In vitro frameshifting assays

Frameshifting levels for SARS-CoV-2 were determined using the p2luc bicistronic dual-luciferase reporter assay system^62^. The SARS-CoV-2 frameshifting sequence (GUUUUUAAACGGGUUUGCGGUGUAAGUGCAGCCCGUCUUACACCGUGCGGCACAGGCACUAGUACUGAUGUCGUAUACAGGGCUUUUGAU) was inserted into the p2luc vector via the BamHI and SacI restriction sites, such that it was located between the Renilla (Rluc) and Firefly (Fluc) luciferases, with Fluc located in the −1 frame relative to Rluc. A control construct was also constructed with a single nucleotide insertion in addition to a disrupted slippery sequence, that causes Fluc to be in the 0 frame relative to Rluc and allows for the normalization of frameshifting readings to 100% (GUUAUUCAAGCGGGUUUGCGGUGUAAGUGCAGCCCGUCUUACACCGUGCGGCACAGGCACUAGUACUGAUGUCGUAUACAGGGCUUUUGAU). The HIV frameshifting element was used as a control (Slippery, UUUUUUAGGGAAGAUCUGGCCUUCCCACAAGGGAAGGCCAGGGA; Control, CUUCUUAAGGGAAGAUCUGGCCUUCCCACAAGGGAAGGCCAGGGA).

mRNA was transcribed using the mMESSAGE mMACHINE T7 Transcription Kit (Invitrogen) and purified using the MEGAclear Transcription Clean-Up Kit, following manufacturer′s protocols. 200 ng of mRNA was mixed with varying concentrations of LNA for 1 hour at room temperature. *In vitro* translation assays were run on the prepared mRNA templates using Rabbit Reticulocyte Lysate kits (Promega), with translation mixes prepared in triplicate as per manufacturer′s protocols (for S2D, all other LNAs are awaiting replicates), followed by incubating at 30°C for 1.5 h (for S2D, all other LNAs will be repeated with this incubation step, however equivalent experiments with S2D without incubation showed no difference in signal). Luciferase readings were performed using the Dual Luciferase Reporter Assay System (Promega) on a Biotek Synergy Neo2 plate reader, using 2 μL of lysate x 3 triplicate wells for each prepared translation mix.

Frameshifting levels were calculated by taking the ratio:

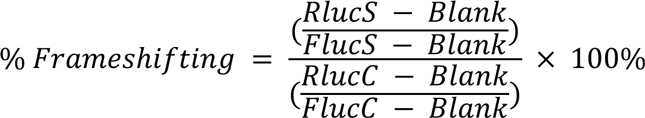

The resulting mean fold-change frameshifting data were then fitted using the Matlab 2019b (MathWorks) “Levenberg-Marquardt” algorithm to a standard binding isotherm (Slp2 was fitted using a constrained IC_50_ to 130 nM):

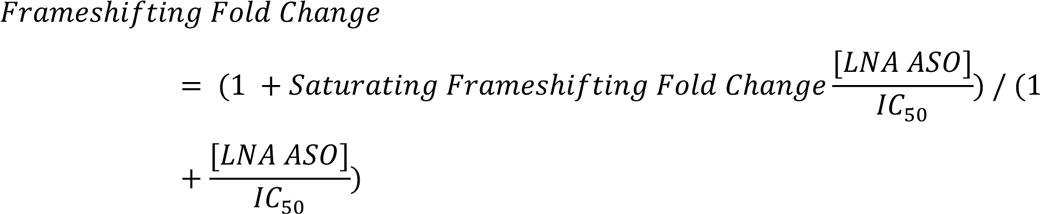

### LNA treatment, caspase activation and cell viability assays

Cell-lines used for *in cellulo* experiments, Vero E6 and Huh-7, were maintained at 37°C in Dulbecco’s modified Eagle medium (DMEM; Gibco) containing 10% fetal bovine serum (FBS; Invitrogen), penicillin and streptomycin (Gibco), and HEPES buffer (Gibco).

For caspase activation and cell viability assays, one day prior to transfection, 10,000 cells per well of Huh-7 or Vero E6 cells were plated in growth media in black-walled, clear bottom 96-well plates (EK-25090, E&K Scientific) to be 50-70% confluent at time of transfection. 24 h later, LNA-modified ASOs were diluted in Opti-MEM I and mixed with Lipofectamine 3000^®^ reagent (Life Technologies) according to manufacturer′s protocol to achieve final LNA concentrations of 1 μM, 0.1 μM, and 0.01 μM, in duplicate. The complexes were incubated at room-temperature for 30 mins, after which 10 μL of each of the LNA-Lipofectamine complex was added dropwise to cells containing 90 μL of growth media. Cells were then incubated as described. A caspase-activating LNA, Pos_LNA, and known apoptotic activator, camptothecin, were used as positive controls.

To test caspase activation, 24 h post-transfection with the LNA ASOs, the Caspase-Glo 3/7 reagent (G8092/G8093 kit; Promega, Madison WI, USA) was prepared at room-temperature according to the manufacturer′s specifications. Once the cell plates were equilibrated at room temperature, a 1:1 ratio of Caspase-Glo 3/7 reagent volume to sample volume was added, its contents were mixed gently at 350–500 rpm for 30 s using a plate shaker, and left to incubate at room temperature for 45 mins in the dark. The plates were measured for caspase activation by luminescence using a Tecan Infinite® multimode plate reader (M1000, Tecan). Data were analyzed and graphed in Prism 8 by GraphPad.

Cell viability after LNA ASO treatment was determined by PrestoBlue cell viability reagent (Invitrogen) following the manufacturer′s instructions. LNA ASO transfections were generated using the same transfection protocol described above in 96-well plates. After 4 days of incubation with the LNA ASO transfection mix, 10 μl Prestoblue reagent (catalog no. 11644807; Roche Applied Science) was added to each well and the plate was incubated for 1-2 h. The plate was then read for fluorescence (540 nm excitation/590 nm emissions) using a Tecan Infinite® multimode plate reader (M1000, Tecan). The cell viability after 4-days of LNA ASO treatment was calculated as a percentage of total cellular viability, normalized to the average viability count of non-treated wells in the absence of LNA ASOs after background correction. Data were analyzed and graphed in Prism 8 by GraphPad.

### Cellular SARS-CoV-2 replication assays

For cellular replication assays, LNA ASOs were reconstituted in RNase-free water at 100 μM stock solutions, aliquoted and stored at −20°C prior to single-use. One day prior to transfection, Huh-7 or Vero E6 cells were plated in 96-well clear bottom plates to be 50-60% confluency at the time of treatment with LNAs S2D, S3D-1, S3D-2, or Scrambled LNA. Lipofectamine 3000® (Life Technologies) was used to transfect LNA ASO’s into cells at 25 nM final concentration per manufacturer′s protocol. Cells were then infected with SARS-CoV-2 reporter virus expressing nanoluciferase (SARS-CoV-2 nLUC) at an MOI of 0.3 for 1 h after which the virus was removed and fresh medium was added. Recombinant SARS-CoV-2 nLUC is an authentic fully replicating virus where ORF7 has been deleted and replaced with nLUC. Thus the measurement of nLUC expression is a surrogate marker of virus replication enabling the screening of antiviral compounds, obviating the need for costly and time-consuming virus titering assays^63^. A nucleoside analog NHC (a.k.a. EIDD-1931) with potent activity against SARS-CoV-2 was included as a positive control^33^. A DMSO control was included as a mock treated, negative control. Data was graphed and analyzed in Prism 8 by GraphPad. Statistical analysis for data from each cell-type was computed as an ordinary one-way ANOVA using Dunnet′s multiple comparisons test against the DMSO control from each cell type.

### Sample preparation for chemical mapping and cryo-electron microscopy

The SARS-CoV-2 FSE sequence was derived from the NCBI GenBank reference sequence NC.045512.2^64^ and corroborated using manual alignment against existing annotated betacoronavirus FSEs^3^. For cryo-EM studies, the sequence was then prepended with the Φ2.5 T7 RNA polymerase promoter (to avoid the obligate 5′ GG coding requirement of the canonical Φ6.5 T7 RNAP promoter)^65^. Primers were designed for PCR assembly using the Primerize^65, 66^ web server, ordered from Integrated DNA Technologies, and assembled into full-length double-stranded DNA (dsDNA) by PCR assembly following the Primerize protocol using Phusion polymerase, ′High-Fidelity′ buffer (Thermo Scientific™), and an annealing temperature of 64°C. Product homogeneity was assessed by 1xTBE (Ambion) – 4% agarose gel electrophoresis as visualized with ethidium bromide. The resulting product was purified using Qiagen QIAquick PCR purification kits, eluted in molecular biology grade double-distilled H_2_O (ddH_2_O, Invitrogen™), and quantified using a NanoDrop spectrophotometer (Thermo Scientific™). The resulting dsDNA was then used for *in vitro* transcription (IVT) using a Thermo Scientific TranscriptAid T7 kit (Thermo Scientific™) where 22 reactions were carried out at 1.19 μg dsDNA per reaction and left at 37°C for 12 hours. The resulting product was split equally over four Zymo25 RNA Clean and Concentrator (Zymo Research) columns and cleaned following the manufacturer′s protocol with an extra 15-minute drying step prior to elution in 30 μL of ddH_2_O per column; RNA was quantified at 5-fold dilution via NanoDrop. Next, the RNA was purified by denaturing Poly-Acrylamide Gel Electrophoresis (dPAGE), using 15% 29:1 Bis-Acrylamide, 7M urea, 1x TBE gels cast in BioRad Criterion 1mm Midi Cassettes that were pre-run at 25 Watts for 1 h, before loading the 90°C denatured RNA in 70% formamide, 1x TBE loading buffer with Bromophenol Blue and Xylene Cyanol running markers and run at 25 Watts for another 45 mins. 50 μL of RNA was loaded per gel, so two gels were used. The side wells of each gel were used for size standards and 2/35ths diluted RNA for purification corroboration. To prevent potentially disrupting RNA structure with intercalating dyes, the major RNA products on the dPAGE gels were localized and excised by brief UV shadowing^67^. After the bulk RNA was excised, the remaining gel was stained for 20 mins in SyBrGold (Invitrogen™) and visualized by UV transillumination. The excised gel pieces were crushed and soaked overnight in 300 μL 10mM Tris, 1mM EDTA buffer (Ambion) per gel slice on an active vortexer at 4°C. The resulting gel slurry was then spin filtered through 0.45 μm filters (Corning® Costar®) and the filtrate was first purified in one Zymo25 RNA Clean and Concentrator column per excised band, eluting in 40 μL ddH_2_O, and then through one Zymo5 RNA Clean and Concentrator column per sample, eluting in 25 μL ddH_2_O, at which point samples were consolidated and concentrated one last time in one Zymo5 RNA Clean and Concentrator column per sample, eluting in 14 μL ddH_2_O. RNA was quantified by NanoDrop. The same overall protocol was carried out for the FSE and FSE-ATP-TTR3 chemical mapping constructs (see below). Chemical mapping constructs were prepended with the canonical Φ6.5 T7 RNAP promoter as the 5′ reference hairpins used had 5′ terminal GGs; additionally, these constructs were gel extracted using blue-light transillumination after staining with SyBrGold. The FSE 1D chemical mapping construct′s assembled dsDNA was 2% agarose gel purified by size using blue-light transillumination using SyBrSafe (Invitrogen™) and purified using a Qiagen MinElute Gel Extraction kit prior to IVT.

### Cryo-EM data acquisition

The RNA samples were folded prior to plunge freezing as previously described^7^. Briefly, the RNA sample in a buffer containing 50 mM Na-HEPES (pH 8.0) was denatured at 90°C for 3 mins and cooled at room temperature (RT) for 10 mins; MgCl_2_ was then added to achieve a final concentration of 15 μM RNA and 10 mM MgCl_2_, and samples were incubated at 50°C for 20 mins, followed by cooling at RT for 3 mins. Three microliters of the FSE and FSE-ATP-TTR3 samples were applied onto glow-discharged 200-mesh R2/1 Quantifoil grids. The grids were blotted for 4 s and rapidly cryocooled in liquid ethane using a Vitrobot Mark IV (Thermo Fisher Scientific) at 4°C and 100% humidity. The final datasets in this study were imaged in a Titan Krios cryo-electron microscope (Thermo Fisher Scientific) operated at 300 kV with GIF energy filter (Gatan) at a magnification of 165,000× (corresponding to a calibrated sampling of 0.82 Å per pixel) for both the samples. Micrographs were recorded by EPU software (Thermo Fisher Scientific) with a Gatan K2 Summit direct electron detector, where each image was composed of 30 individual frames with an exposure time of 6 s and an exposure rate of 8.3 electrons per second per Å^2^. A total of 10,222 movie stacks for FSE and 12,380 movie stacks for FSE-ATP-TTR3 were collected.

### Single-particle image processing and 3D reconstruction

All micrographs were first imported into Relion for image processing. The motion-correction was performed using MotionCor2^68^ and the contrast transfer function (CTF) was determined using CTFFIND4^69^. All particles were autopicked using the NeuralNet option in EMAN2. Then, particle coordinates were imported to Relion, where the poor 2D class averages were removed by several rounds of 2D classification. The initial models for both datasets were built in cryoSPARC using the ab-initio reconstruction option. For the FSE, 1,063,711 particles were picked and 445,707 were selected after 2D classification in Relion. After removing classes with poorly connected density by 3D classification, the final 3D refinement was performed using 109,137 particles in Relion, and a 6.9-Å map was obtained. For the FSE-ATP-TTR3, 1,103,091 particles were picked and 902,309 were selected after 2D classification in Relion. After removing classes with poorly connected density by 3D classification in Relion, two rounds of heterogeneous refinement performed in cryoSPARC to further remove contaminant particles. The final 3D homogenous refinement was performed using 257,558 particles, and a 6.4-Å map was obtained. Resolution for the final maps was estimated with the 0.143 criterion of the Fourier shell correlation curve without or with mask. A Gaussian low-pass filter was applied to the final 3D maps displayed in the UCSF Chimera software package^70^.

### One-dimensional chemical mapping

Chemical mapping was conducted on FSE, plusFSE and FSE-ATP-TTR3 constructs (**Extended Data Table 2**) with reference hairpins added to either side of the regions of interest^71^. In brief, dPAGE purified RNA was diluted to 80 nM in 50 mM Na-HEPES at pH 8.0, 10 mM MgCl_2_ folded using the same folding scheme described for cryo-EM (plus an extra incubation at 37°C for 24 mins at the end), followed by dilution to 60 nM in either 0.5% Dimethyl Sulfate (DMS) in 0.5% ethanol in ddH_2_O, or 4.24 mg/mL 1-methyl-7-nitroisatoic anhydride (1M7) in Dimethyl Sulfoxide (DMSO), reaction at room temperature for 15 mins, followed by quenching in 500mM NaMES at pH6.5 for 1M7, and 51% β-mercaptoethanol (β-ME) respectively. During this quench step, FAM-labelled Tail2 reverse transcription primer was added to a final concentration of 2 nM, with the RNA at 40 nM. Reverse transcription was then carried out using SuperScript III (Invitrogen™) alongside an RNA dideoxynucleotide (ddNTP) sequencing ladder. The resulting cDNA was cleaned and then resuspended in HiDi-formamide (Applied Biosystems™) with ROX350 (Applied Biosystems™) added as fiducial markers. Concentrated and dilute samples were then submitted for capillary electrophoresis at Elim Biopharmaceuticals and the resulting FAM channel traces, as aligned by the ROX350 markers, were used in HiTRACE^71^ to assign per-nucleotide reactivity data for DMS, 1M7, and negative control incubation conditions. The resulting .rdat files were then processed using *RNAStructure* with 100 bootstrapping rounds, allowing for the computation of pseudoknots with the Biers software (https://github.com/ribokit/Biers).

### Two-dimensional chemical mapping (M2-seq)

Following one-dimensional chemical mapping, the same constructs were also subjected to two-dimensional (2D) chemical mapping which uses DMS modification on folded RNA but leverages information on structural perturbations from sequence mutations to more directly infer helices^72^. The resulting signals are read out using Illumina short read sequencing, exploiting the mutational readthrough of DMS modified bases by the retrotranscriptase TGIRT-III (InGex).

In brief, the same DNA encoding the 1D chemical mapping constructs were first subjected to error-prone PCR (epPCR) where 2 ng/μL of PCR assembled (see above) dsDNA (1.6μg total dsDNA used per construct, 8⨉ epPCRs) was amplified using the epPCR forward and reverse primers at 100 μM (Extended Data Table 2) under error-prone conditions: 10 mM Tris-HCl pH 8.3, 50 mM KCl, 7 mM MgCl_2_, 1 mM dTTP, 1 mM dCTP, 200 nM dATP, 200 nM dGTP, 500 nM MnCl_2_, and 1x Taq polymerase (New England BioLabs inc.), using the following PCR cycling conditions: initial denaturation at 98°C for 60 s; then 35 cycles of 94°C for 30 s, 60°C for 60 s, and 72°C for 180 s; followed by 72°C for 10 mins. The resulting PCR products were concentrated using Qiagen QIAquick PCR purification kits (1 per 200μL product). The resulting cleaned dsDNA was purified by size on a SyBrSafe-stained 2% agarose gel using blue-light transillumination and purified using a Qiagen MinElute Gel Extraction kit prior to IVT (4x TranscriptAid reactions, 2000ng dsDNA per reaction, 12 h reaction at 37°C followed by 20 mins DNase I treatment at 37°C, 1 μL per reaction volume). The resulting RNA was Zymo25 purified, and then 8% dPAGE purified (see above) by size using SyBrGold, blue-light transillumination, and one ZR small-RNA PAGE Recovery kit per construct following the manufacturer′s protocol. The resulting size purified RNA was then subjected to the M2-seq protocol. Briefly, unmodified (nomod) and DMS-modified samples were prepared, using 5 pmol and 12.5 pmol of RNA per condition, respectively. RNA was diluted into 10 mM Tris, 1 mM EDTA buffer (Ambion) to 3.0 μL, denatured at 90°C for 3 mins, cooled to room temperature for 10 mins, and then diluted into 5.0 μL 1.5M sodium-cacodylate pH 7.0, 50 mM MgCl_2_ buffer and 14.5 μL ddH_2_O and allowed to refold at 50°C for 20 mins followed by cooling to room temperature for 3 mins. Following this folding, the buffered RNA was then subjected to DMS chemical modification in 2.5 μL 15% DMS (diluted in 100% ethanol), or just 85% ethanol for the no modification condition, at 37°C for 6 mins, followed by quenching with 25.0 μL β-mercaptoethanol. All of the DMS and β-mercaptoethanol steps were carried out in a fume hood. The resulting quenched RNA reactions were then diluted in ddH_2_O to 100 μL and ethanol precipitated using GlycoBlue (Invitrogen™) as a co-precipitant, resuspending the resulting RNA in 7.0 μL ddH_2_O. Next, the cleaned nomod and DMS-modified RNA was reverse transcribed using RTB FAM-labelled primers (**Extended Data Table 2**) using a unique index per condition, per construct. 4.6 μL of RNA was incubated with 0.930 μL of 285 nM RTB primer in 6.520 μL retrotranscription buffer: 2.4 μL 5x TGIRT-III buffer, 1.2 μL 10mM dNTPs mix (New England BioLabs), 0.6 μL 100mM DDT (Invitrogen™), 1.32 μL ddH_2_O, and 1.0 μL TGIRT-III enzyme. The reactions were allowed to sit at 25°C for 5 mins, before being kept at 64°C for 3 h. Afterwards, the reactions were treated with 5.0 μL 400 mM NaOH at 90°C for 3 mins, acid quenched (see above) with 2.2 μL quench acid, and diluted in 30.75 μL ddH_2_O prior to purification using a Zymo Oligo Clean and Concentrator kit, eluting in 12.5 μL ddH_2_O. The resulting cleaned cDNA were then individually subjected to unidirectional PCR for second strand synthesis using the second strand synthesis primer (**Extended Data Table 2**). 1.0 μL 100 μM primer was used per 2.5 μL cDNA, and 19.5 μL PCR master mix (same as PCR assembly, see above). The reaction mixtures were initially denatured at 98°C for 60 s; then 4x cycles of 98°C for 10 s, 58°C for 30 s, and 72°C for 30 s; followed by 72°C for 10 mins. Immediately after the final step, PCR amplification using the iTru p5 and p7 primers (**Extended Data Table 2**) was conducted in the same reaction mixtures. 2.0 μL of equimolar p5/p7 (50 μM per primer) mixture was spiked into each uni-directional PCR reaction product, mixed, spun down, and quickly returned to the thermal cycler to conduct: initial denaturation at 98°C for 60 s; then 5x cycles of 98°C for 10 s, 50°C for 30 s, and 72°C for 30 s; followed by 15x cycles of 98°C for 10 s, 60°C for 30 s, increasing this temperature incrementally each cycle to a final temperature of 70°C by cycle 15, and 72°C for 30 s; then 72°C for 10 mins. The resulting reaction products were then concentrated using Qiagen QIAquick PCR purification kits, and finally 1% agarose gel purified by size using blue-light transillumination using SyBrSafe and purified using Qiagen MinElute Gel Extraction kits. The resulting dsDNA sequencing libraries were then sequenced using an Illumina MiSeq using the v3 600 cycle kit, loading equimolar amounts of DMS treated DNA per construct, and 3 fold less nomod DNA per construct (14 fmol dsDNA total including 40% PhiX control).

M2-seq data analysis was performed using the pipeline at https://github.com/ribokit/M2seq. FASTQ files were demultiplexed with NovoBarcode, and demultiplexed files were processed with ShapeMapper to obtain mutation strings for each read in a binary format. The simple_to_rdat.py script was used to generate 2D mutational profiles in RDAT files. These files were then analyzed with Biers (https://ribokit.github.io/Biers/) to generate Z-scores, normalized 1D DMS profiles, and secondary structure predictions using ShapeKnots with 100 bootstrapping iterations guided by both Z-scores and 1D DMS profiles.

### Reactivity-guided structure modeling for extended FSE

Secondary structure modeling was performed for FSE constructs including 110 upstream nucleotides, guided by SHAPE reactivity data collected in this study along with various recently published SHAPE and DMS reactivity datasets^27, 28, 73^. SHAPE reactivity data were collected for the extended FSE constructs with 1M7 as described above; data were averaged across two replicates. DMS reactivity data from^27^ were normalized by scaling the median of the top 5% of reactive bases to 1.0 and winsorizing. Predictions reported here were made with ShapeKnots and 100 bootstrapping iterations if pseudoknots were predicted; otherwise, RNAStructure′s Fold algorithm was used, again with 100 bootstrapping iterations. All secondary structures were depicted using RiboDraw (https://github.com/ribokit/RiboDraw/).

### FSE model building

The 6.9-Å cryo-EM FSE map was first subjected to segmentation using Segger^48^ to remove any contours lower than 0. The resulting segmented map was then low-pass filtered to 20-Å by autoDRRAFTER, and a map threshold of 0.05 was used to compute the autoDRRAFTER fitting nodes. AutoDRRAFTER identifies graph end nodes in both the experimental cryo-EM map and the secondary structure, and then seeks to match these end nodes together to initiate model building. Since the FSE is expected to have only one stem-loop (one end node), the SARS-CoV-2 FSE Stem 3^21^, Stem 3 was hence used by the software as the initial point for docking the secondary structure into the cryo-EM map. AutoDRRAFTER identified two map end-nodes (**Extended Data Fig. 7j**) into which this Stem 3 was automatically docked and both were used for two identical, and converged runs using Stem 3 as the docked helix, using the literature secondary structure as a constraint (see **Extended Data Table 2**). We noted that jobs explicitly initiated with Stem 3 docking into what was later determined to be the incorrect region of the density (using the FSE-ATP-TTR-3) produced results very similar to jobs initiated with the other, correct, node used as the docking point (**Extended Data Fig. 7k**). After this set of modelling as well as the cross-validation using FSE-ATP-TTR-3, subsequent autoDRRAFTER jobs were conducted only using the correct node as the specified docking position for Stem 3. Ultimately, jobs run using the 1D chemical mapping-derived secondary structure resulted in marginally (∼0.8 Å lower estimated accuracy) better correlated models compared to models generated with either the literature and M2-seq secondary structures (**Extended Data Fig. 7a-i**) and so were used for subsequent analysis. Regardless of the starting secondary structure, all autoDRRAFTER runs resulted in similar global tertiary structures (**Extended Data Fig. 7a-i**). Modelling was attempted using the FSE-ATP-TTR3 map, using the previously computed ATP-TTR 3 model as the initial docking structure, but the absence of significant density at the FSE portion of the map produced convergent results that only confidently placed the Stem 3 helix (**Fig. 3c**). All jobs were run on the Stanford high performance computing cluster, Sherlock 2.0, using the latest distribution of Rosetta and autoDRRAFTER.

### FSE-ATP-TTR3 design

ATP-TTR-3, a stabilized aptamer for ATP and AMP, was selected as a tag due to its stability, structural features conducive to projection angle identification, and its helical end for linkage. The tag was linked to the FSE by removing the dimerization loop (44-52) and placing the tag in its place. Nucleotides were inserted in the FSE-tag junction (C44G and U52A, FSE numbering) to remove any predicted secondary structure interactions between the FSE and tag and retain the individual secondary structure prediction for FSE and the ATP-TTR 3 tag. Secondary structure was predicted using CONTRAfold 2.0^74^ using the following command: contrafold predict seq.fasta.

### FSE-ATP-TTR3 Density Fitting

The previously obtained density of ATP-TTR 3 and the FSE density were fit into the density obtained for the FSE-ATP-TTR3 construct using Situs ^49^. The overlap was not expected to be ideal because the FSE-ATP-TTR3 construct does not contain the dimerization loop of the FSE (nts 44-52) and has extra bases, C44 and U175, inserted at the fusion interface between the FSE and ATP-TTR 3, a linkage not present in either individual construct. First, the ATP-TTR 3 density was fit into the FSE-ATP-TTR3 density using the commands:

> vol2pdb ATP-TTR_apo_ribosolve.mrc atp.pdb
>
> colores FSE-ATP-TTR3.mrc atp.pdb -res 4

Next, the docking result with the highest correlation was subtracted from the FSE-ATP-TTR3 density.

> pdb2vol atp_best_001.pdb atp_best_001.mrc #using 1A voxel spacing
>
> voldiff FSE-ATP-TTR3.mrc atp_best_001.mrc diff.mrc

Finally, the FSE density was fit into the difference density.

> vol2pdb FSE-wt.mrc fse.pdb
>
> colores diff.mrc fse.pdb -res 7
>
> pdb2vol col_best_001.pdb fse_001.mrc #same for all 9 hits, using 1A voxel spacing

The results were visualized in Chimera. The stem attached to the ATP-TTR 3 on the docking result with the highest correlation was assigned to stem 3.

### De novo RNA 3D structure modeling

*De novo* models were generated for frameshift stimulation elements from seven betacoronaviruses with Rosetta′s *rna_denovo* application using the standard FARFAR2 protocol^75^. Secondary structures for the frameshift stimulation elements for murine hepatitis virus (MHV), human coronavirus HKU1, human coronavirus OC43, bovine coronavirus (BCoV), Middle Eastern respiratory syndrome-related coronavirus (MERS), and SARS coronavirus (SARS-CoV-1) were obtained from prior literature analyses^21^. For MERS, 2,000 decoys were produced using the FARFAR2 ROSIE server. *De novo* models for the SARS-CoV-2 FSE models were obtained as described previously^44^. For all other cases, 150,000 decoys were generated using the Stanford high performance computing cluster, Sherlock 2.0, and the Open Science Grid^76^. In each case, the 400 top-scoring models were clustered using the rna_cluster Rosetta application with a 5-Å cluster radius. From the top 10 clusters, top-scoring cluster members demonstrating the threaded and unthreaded topologies were selected. Models are available at https://github.com/DasLab/FARFAR2_FSE_homology.

### phyloP analysis

phyloP scores were extracted directly from the UCSC SARS-CoV-2 genome browser^22^ for both the whole genome and the FSE.

## Acknowledgments

We thank the administrative staff of the Biochemistry Department at Stanford; the SLAC National Accelerator Laboratory; and the Stanford University COVID-19 research oversight committee for supporting conduct of these studies during a university-wide shutdown. We thank Paul Berg for extensive discussion and advice, Corey Hecksel and Patrick Mitchell for expert maintenance of Stanford-SLAC Cryo-EM facilities, Kalli Kappel for advice on using autoDRRAFTER, Helen Berman and Brinda Vallat for RNA model validation, Menashe Elazar for advice on viral replication and cytotoxicity assays, Ved Topkar for advice and assistance with M2-seq, Andrew Watkins for running *de novo* modeling on the Open Science Grid, Adam Siepel and Ritika Ramani for assistance in interpreting the phyloP data, Jody Puglisi for draft review, discussion and advice, and Jennifer Kong and Chandni Patel for providing reagents for PAGE gels. This work was supported by the National Institutes of Health grants (S10OD021600, R01AI148382, P01AI120943, R01GM079429, P41GM103832, U24GM129564 to W.C.; R35GM122579 to R.D; and 1R01AI13219101 to J.S.G.); DOE Office of Science through the National Virtual Biotechnology Laboratory, a consortium of DOE national laboratories focused on response to COVID-19, with funding provided by the Coronavirus CARES Act (W.C. and R.D.); National Science Foundation Graduate Research Fellowship Program grant no. 1650114 (R.R.); NIH/NIDDK 5T32DK007056-44 (E.A.P.); a Stanford ChEM-H Physician Scientist Research Fellowship (E.A.P.); a Stanford ChEM-H COVID-19 Drug and Vaccine Prototyping seed grant (P. Berg, R.D., J.S.G, W.C.); a Stanford ChEM-H Postdocs at the Interface seed grant (R.J.H.); USAMRAA DOD W81XWH1810647 (J.S.G.), Harrington Scholar Innovator Grant (J.S.G.), and Open Philanthropy (J.S.G.); and a Stanford Paul and Mildred Berg Graduate Fellowship (I.Z.).

## Author contributions

W.C., R.D., J.S.G., K.Z. I.Z., and R.J.H. conceived the study and designed the experiments. I.Z. prepared the RNA samples, performed the chemical mapping, and built the models; K.Z. performed cryo-EM sample preparation, screening, data collection, image processing, and structure determination; S.L. assisted with cryo-EM image processing. R.K. designed the FSE-ATP-TTR3 RNA construct and performed the density fitting. I.Z. designed S2D, RL4, and JL4. R.J.H. designed all other LNAs with input from E.A.P.. Y.H. made and titered the recombinant SARS-CoV-2 nLUC virus used in the studies. R.J.H. and T.S. performed *in cellulo* assays. M.W. and R.H. performed *in vitro* frameshifting assays. G.P. and R.K. made movies and carried out cryo-EM correlation analysis. R.R. ran RNAStructure analysis, compiled published data from external labs for assessment, and carried out *de novo* modeling. W.K. conducted the FSE-LNA binding assay with replicates and validation by C.B.S.. W.C., R.D., J.S.G., K.Z., I.Z., R.J.H., R.R., S.L., G.P., M.W., R.H., and V.D. analyzed the data. K.Z., I.Z., R.J.H., W.C., and R.D. wrote and edited the manuscript with input from all other authors.

## Data availability

Cryo-EM maps of the FSE and FSE-ATP-TTR3 have been deposited in the Electron Microscopy Data Bank under accession codes EMD-22296 and EMD-22297; Atomic models of FSE have been deposited in the Protein Data Bank under accession PDB ID code 6XRZ. The raw cryo-EM micrographs are being deposited in EMPIAR.

---

**Extended Data Fig. 1.**
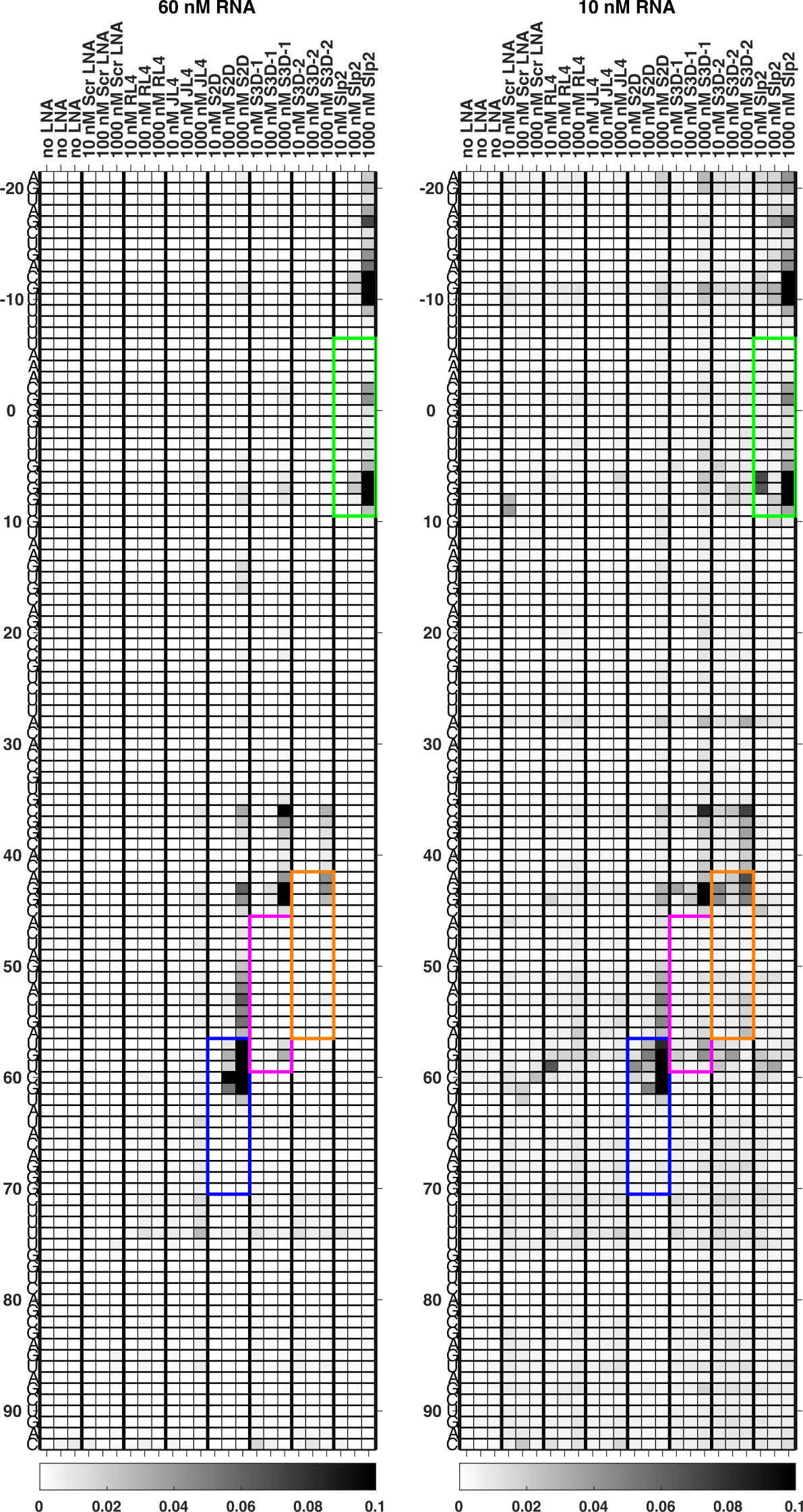
Reverse transcription termination probabilities of SARS-CoV-2 FSE in the presence of LNA ASOs. Data were acquired through capillary electrophoresis of reverse transcription products analyzed in HiTRACE^71,78^ Colored boxes indicate where LNAs are expected to bind based on Watson-Crick complementarity,

**Extended Data Fig. 2.**
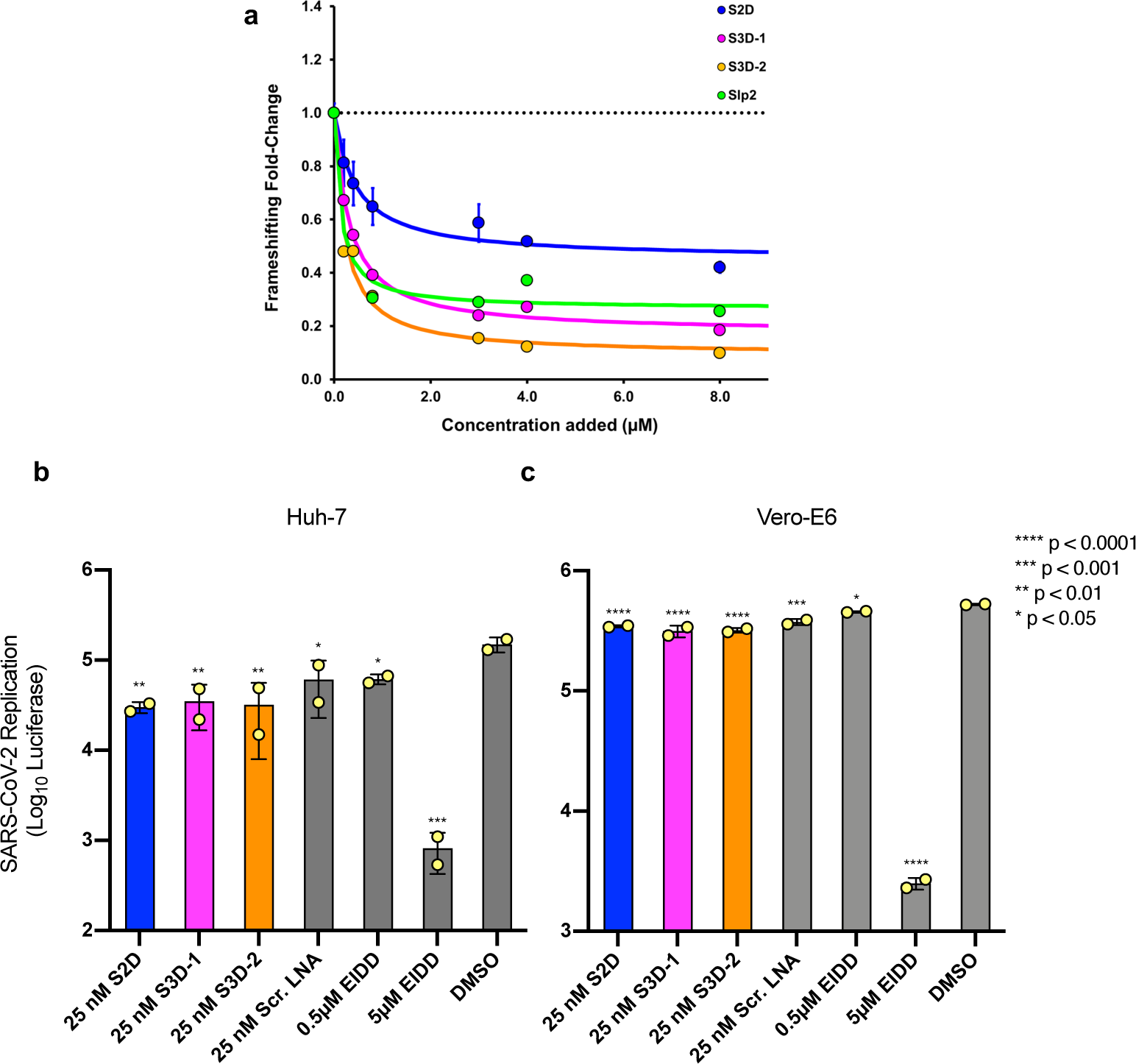
FSE-LNA inhibition *in vitro* dual-luciferase frameshifting assay and of SARS-CoV-2-nLuc virus replication. **a**. Dual-luciferase frameshifting results of FSE-targeting LNA ASOs; same data as main text Fig. 1c, fit to a standard binding isotherm with values of (IC_50_ ± standard error, saturating frameshifting fold change ± standard error; see Methods) as follows: S2D (440 ± 140 nM; 0.45 ± 0.04), S3D-1 (310 ± 35 nM; 0.17 ± 0.02), S3D-2 (210 ± 55 nM; 0.09 ± 0.05), Slp2 (130 ± 130 nM; 0.27 ± 0.04) **b-c**. Viral luciferase reporter data shown on log scale; same data as main text Fig. 1f-g. Cells were pretreated with 25 nM of FSE-directed LNAs or Scr. LNA 24 h prior to infection with SARS-CoV-2-nLuc reporter virus. Luciferase expression was measured 48 h post-infection. The nucleoside analog EIDD-1931 was used as a positive control. Results are shown as percent luciferase expression relative to DMSO control, n = 2. P values were generated by GraphPad Prism software and computed as an ordinary one-way ANOVA using Dunnet′s multiple comparisons test against the DMSO control from each cell type. Error bars represent ± standard deviation.

**Extended Data Fig. 3.**
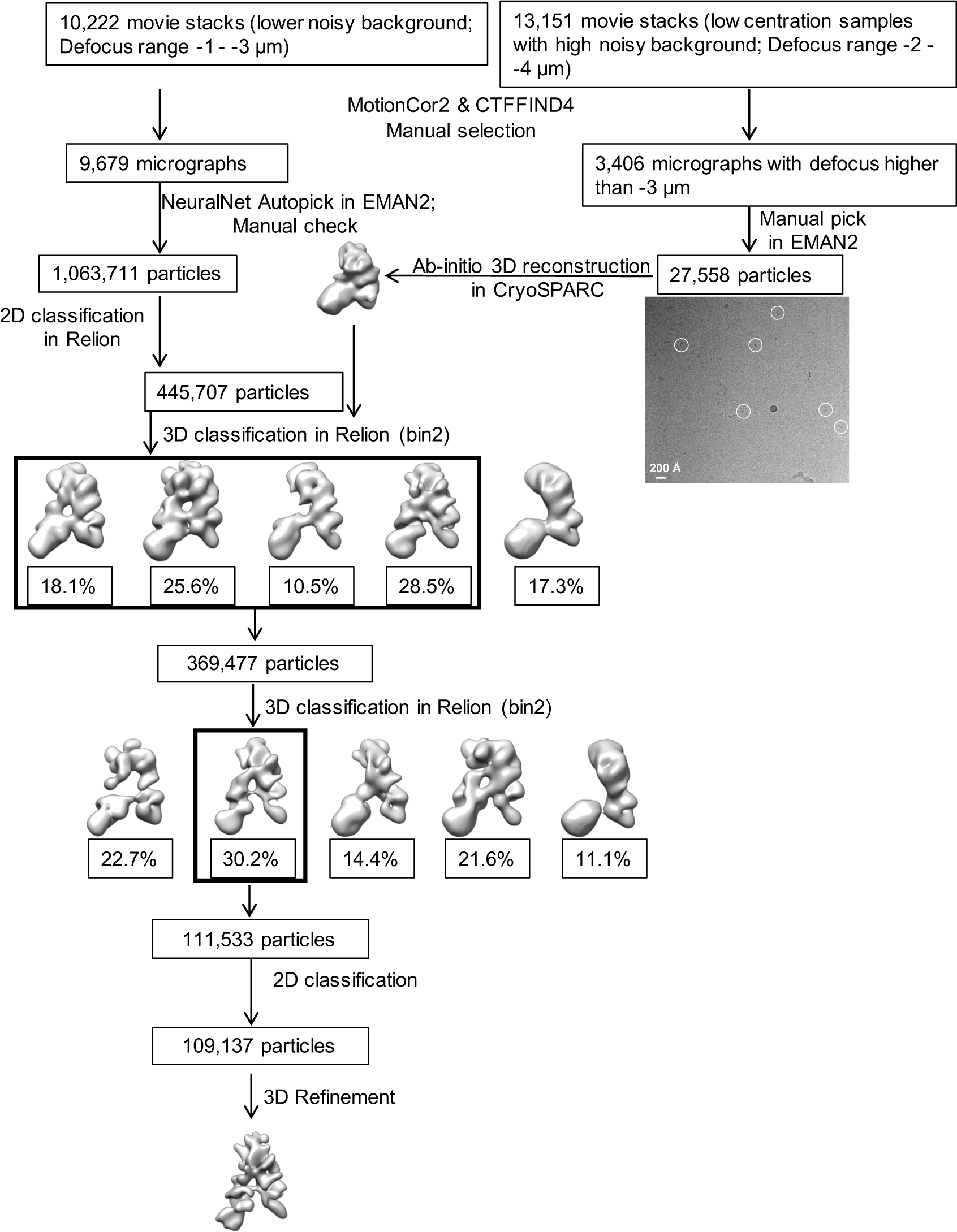
Workflow of cryo-EM data processing of FSE.

**Extended Data Fig. 4.**
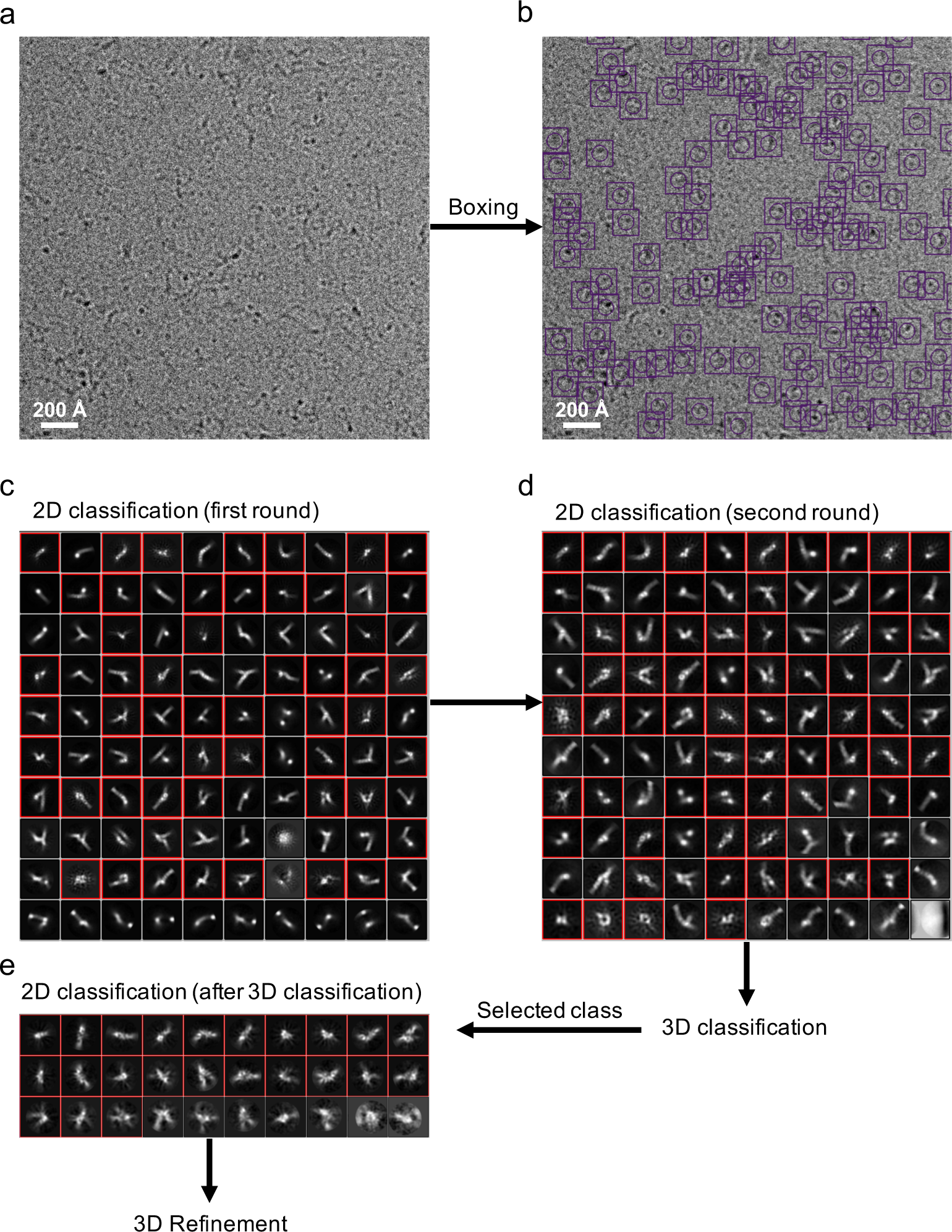
Boxing and 2D classification steps in single-particle cryo-EM analysis of the FSE. **a.** Representative motion-corrected cryo-EM micrograph. **b.** The same micrograph shown in (a) with particles picked by the NeuralNet autopicking option in EMAN2. Circle and square refer to particle size and box size, respectively. **c-e.** Results of reference-free 2D class averages from first and second rounds of 2D classification, and after 3D classification. Classes indicated in red squares were selected for further processing.

**Extended Data Fig. 5.**
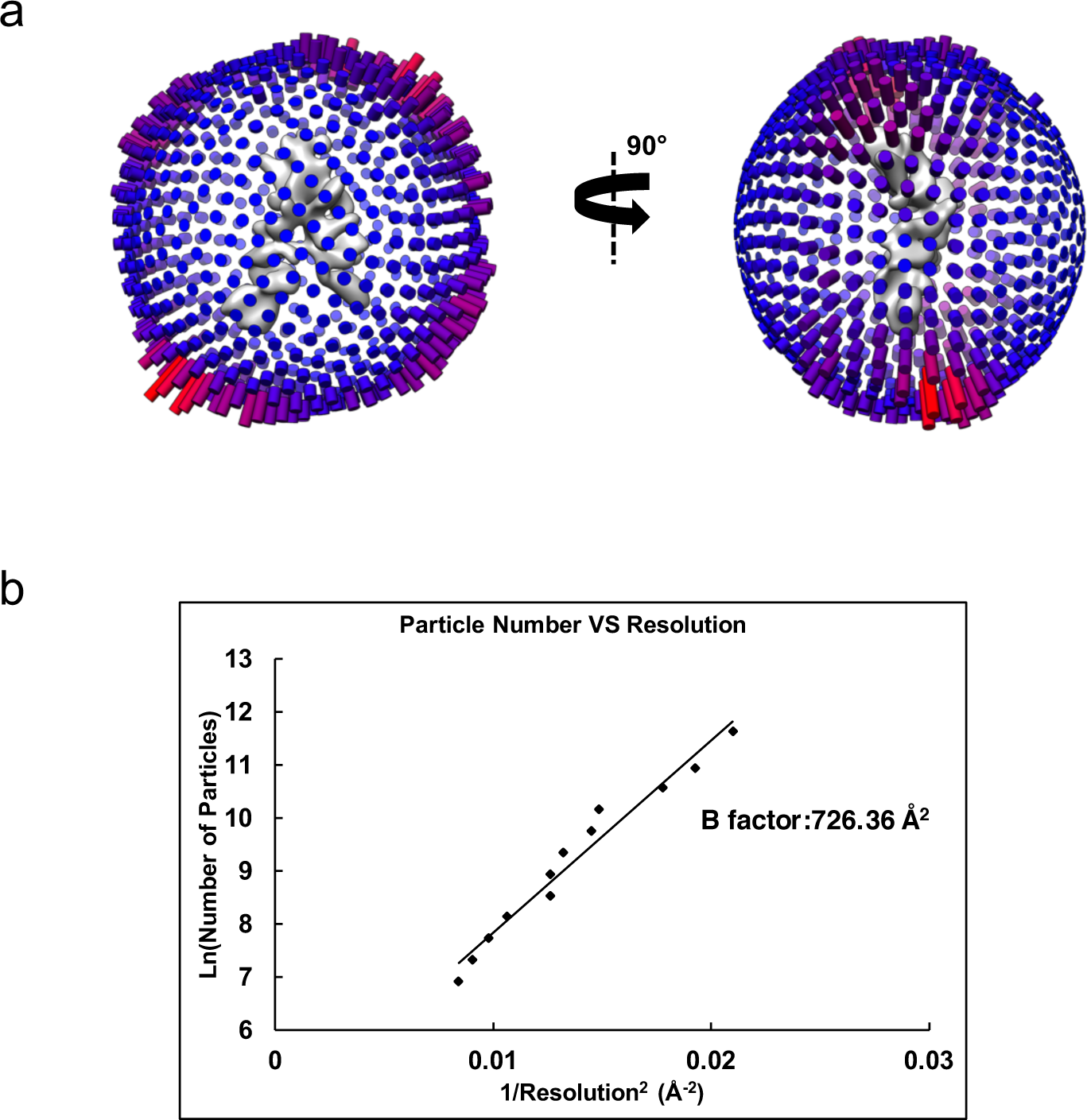
Quality of the particle images in single-particle cryo-EM analysis of the FSE. **a.** Euler angle distribution of the particle images. **b.** Plot of the particle number vs the reciprocal squared resolution. The B-factor was calculated as 2⨉ the linear fitting slope^41,42^.

**Extended Data Fig. 6.**
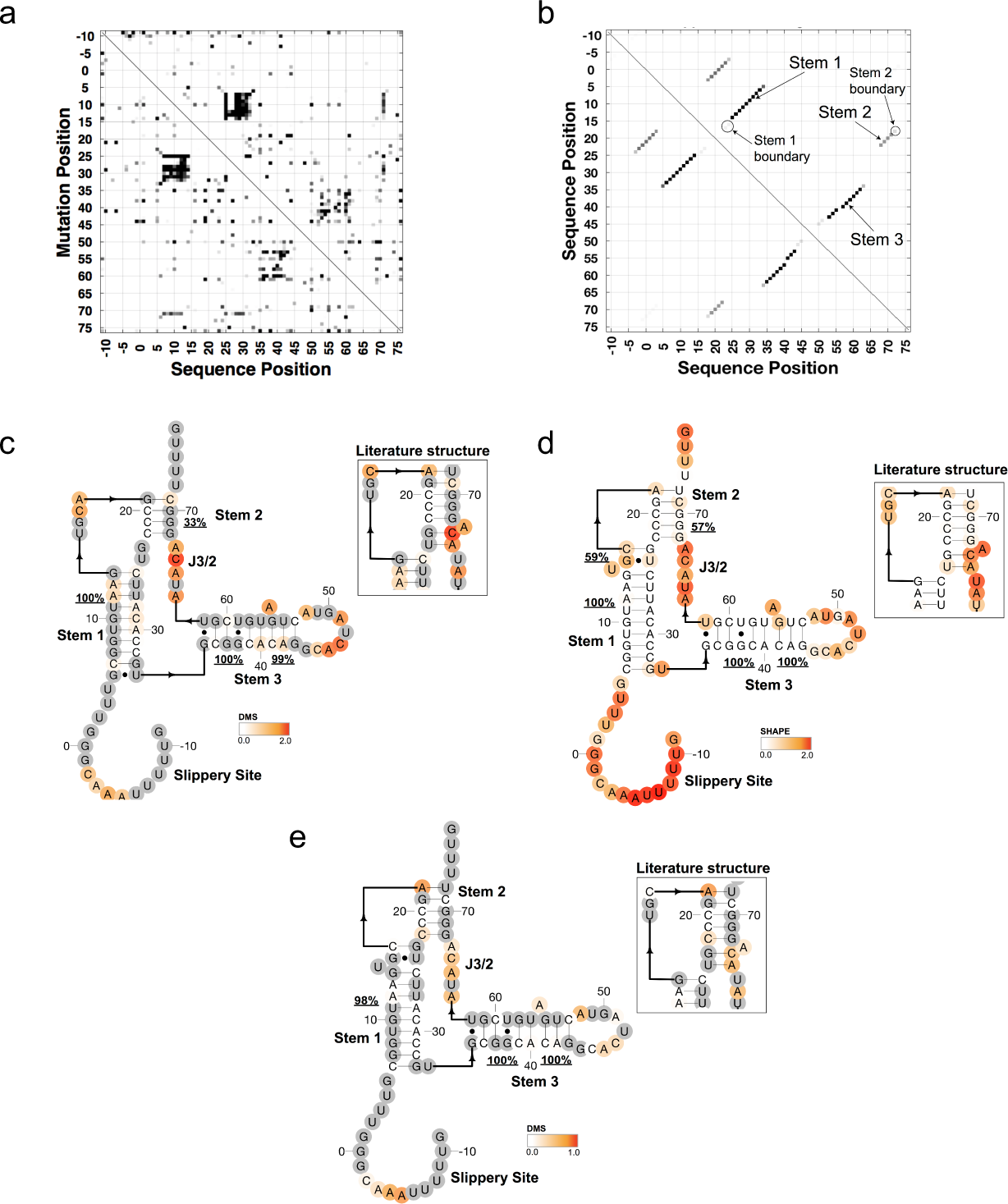
Chemical mapping-derived secondary structures of the SARS-CoV-2 Frameshift Stimulation Element (FSE). **a.** M2-seq Z-score plot for the 88-nt FSE, with numbering convention as used in the main text. **b-c.** Bootstrap confidence values for each base-pair (b) and secondary structure of the FSE (c), as determined by using ShapeKnots guided by M2-seq Z-scores and the 1D DMS chemical mapping signal from M2-seq. Bootstrapping (100 iterations) support for each continuous helix is shown as an underlined percentage. Nucleotides are colored by 1D DMS chemical mapping signal, and nucleotides unreactive to DMS are depicted in grey. Inset depicts secondary structure reported in the literature^3, 4^. **d.** Secondary structure of the FSE as determined by 1D chemical SHAPE mapping with SHAPE reactivity data overlaid. Structure shown here is computed from ShapeKnots allowing for pseudoknots. Bootstrapping (100 iterations) support for each continuous helix is shown as an underlined percentage. **e.** Secondary structure as determined by 1D SHAPE mapping with DMS reactivity data overlaid. Helices that are also predicted with ShapeKnots guided by DMS mapping have bootstrapping support indicated.

**Extended Data Fig. 7.**
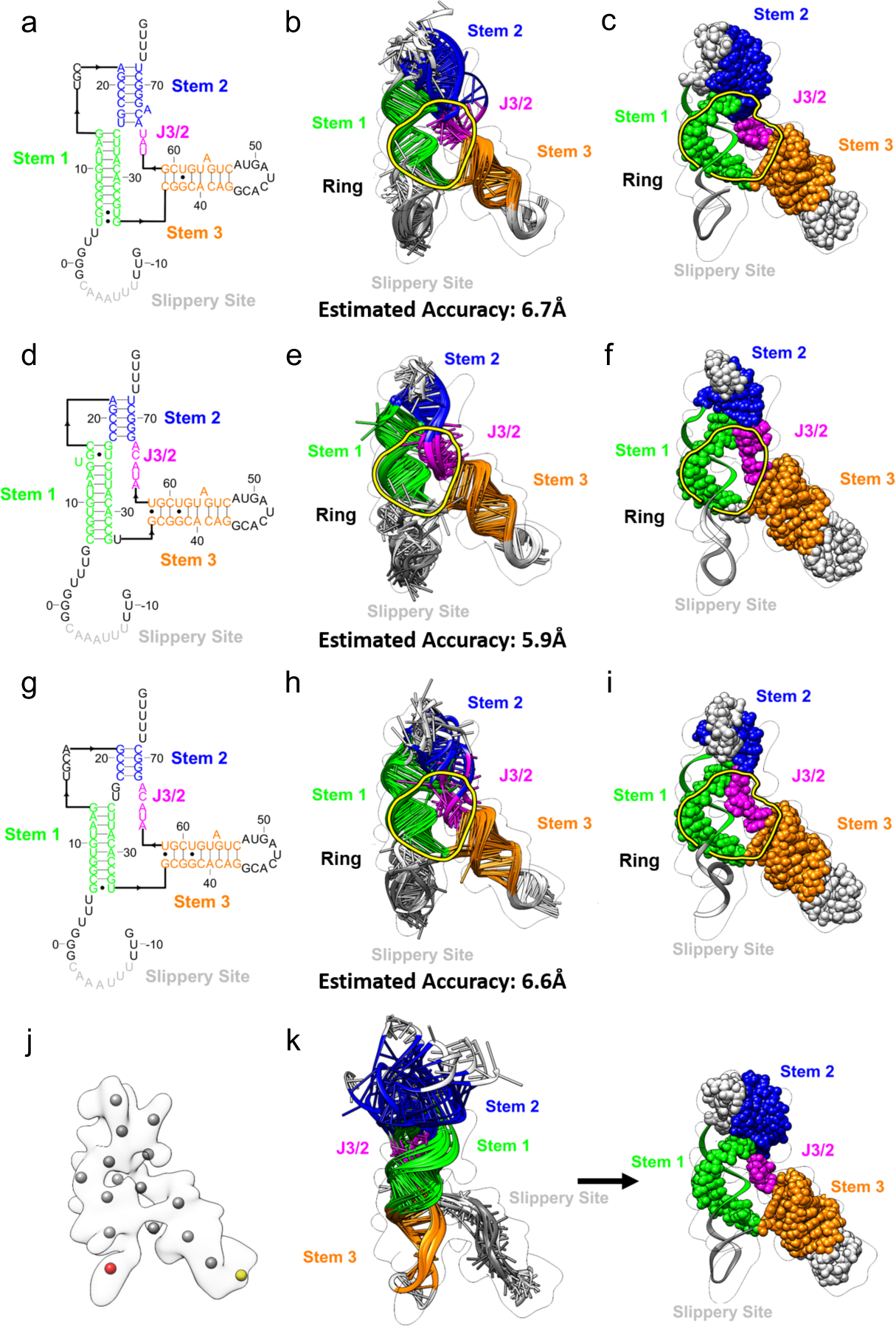
Consistent 5′-threaded tertiary structures of the SARS-CoV-2 FSE using autoDRRAFTER with different modeling assumptions. **a.** The ‘literature’ secondary structure^3, 4^. **b.** Ensemble of the top ten autoDRRAFTER results for (a), with a final mean pairwise root mean squared distance (RMSD) equivalent to an estimated accuracy of 6.7Å^7^. **c.** Simplified view indicating the threading event seen in all top ten results for (a). **d.** The secondary structure as determined by 1D chemical mapping using 1M7 (see Methods) **e.** Ensemble of the top ten autoDRRAFTER results for (d), with a final mean pairwise RMSD equivalent to an estimated accuracy of 5.9Å. **f.** Simplified view indicating the threading event seen in all top ten results for (d). **g.** The secondary structure as determined by 2D chemical mapping using DMS (see Methods) **h.** Ensemble of the top ten autoDRRAFTER results for (g), with a final mean pairwise RMSD equivalent to an estimated accuracy of 6.6Å. **i.** Simplified view indicating the threading event seen in all top ten results for (g). **j-k.** autoDRRAFTER modelling forcing an initial placement of Stem 3 in an alternative position of the map (red vs. yellow in (j)) initially gives poorly converged models that are not well-placed in the density, but after iterative refinement, helices positions shift and lead to well-converged models (k), indistinguishable to results from unbiased autoDRRAFTER modeling (b,e,h).

**Extended Data Fig. 8.**
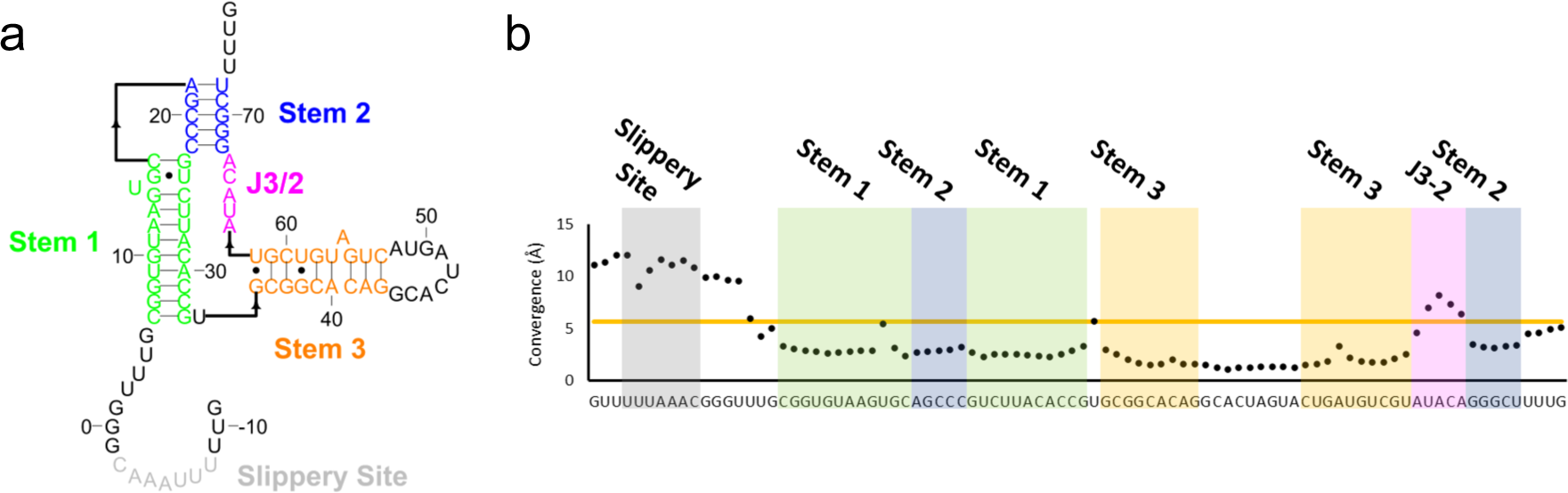
Per-nucleotide modelling convergence of the SARS-CoV-2 Frameshift Stimulation Element (FSE). **a.** The same secondary structure as shown in Fig 2. diagrammed here for ease of reading. **b.** pairwise root mean squared deviation at each nucleotide position over the top 10 autoDRRAFTER derived models, orange line indicates the global mean pairwise root mean squared deviation of 5.68 Å.

**Extended Data Fig. 9.**
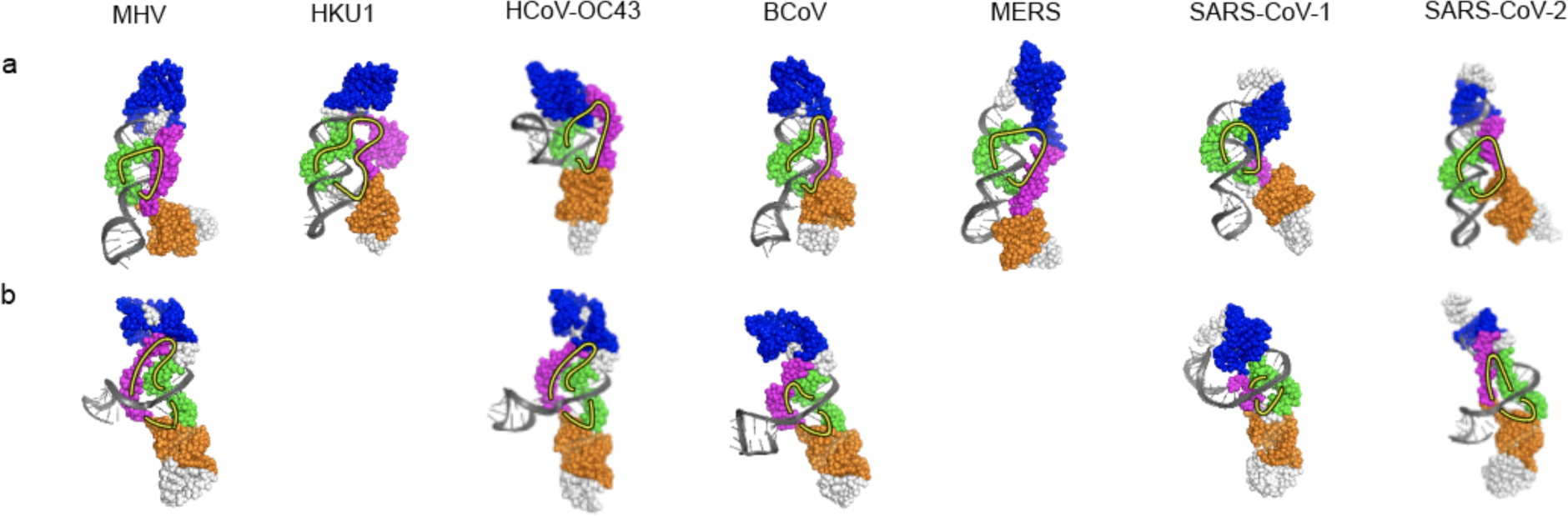
*De novo* models from Rosetta FARFAR2 for Frameshift Stimulation Elements from a range of betacoronaviruses. Murine hepatitis virus (MHV), human coronavirus HKU1 (HKU1), human coronavirus OC43 (HCoV-OC43), bovine coronavirus (BCoV), Middle Eastern respiratory syndrome-related coronavirus (MERS), SARS coronavirus (SARS-CoV-1) and SARS-CoV-2 FSEs computed. Top-scoring cluster centers that show (**a**) 5′-threaded topologies and (**b**) unthreaded topologies are depicted. Models are only shown if a cluster in the top-scoring 10 clusters shows the desired topology. The 5′ strand of Stem 1 and upstream slippery site are colored in dark gray, the 3′ strand of Stem 1 is colored in green, the junction J2/3 is colored in magenta, Stem 2 is colored in blue, and Stem 3 is colored in orange. Yellow circles indicate the hole in each structure.

**Extended Data Fig. 10.**
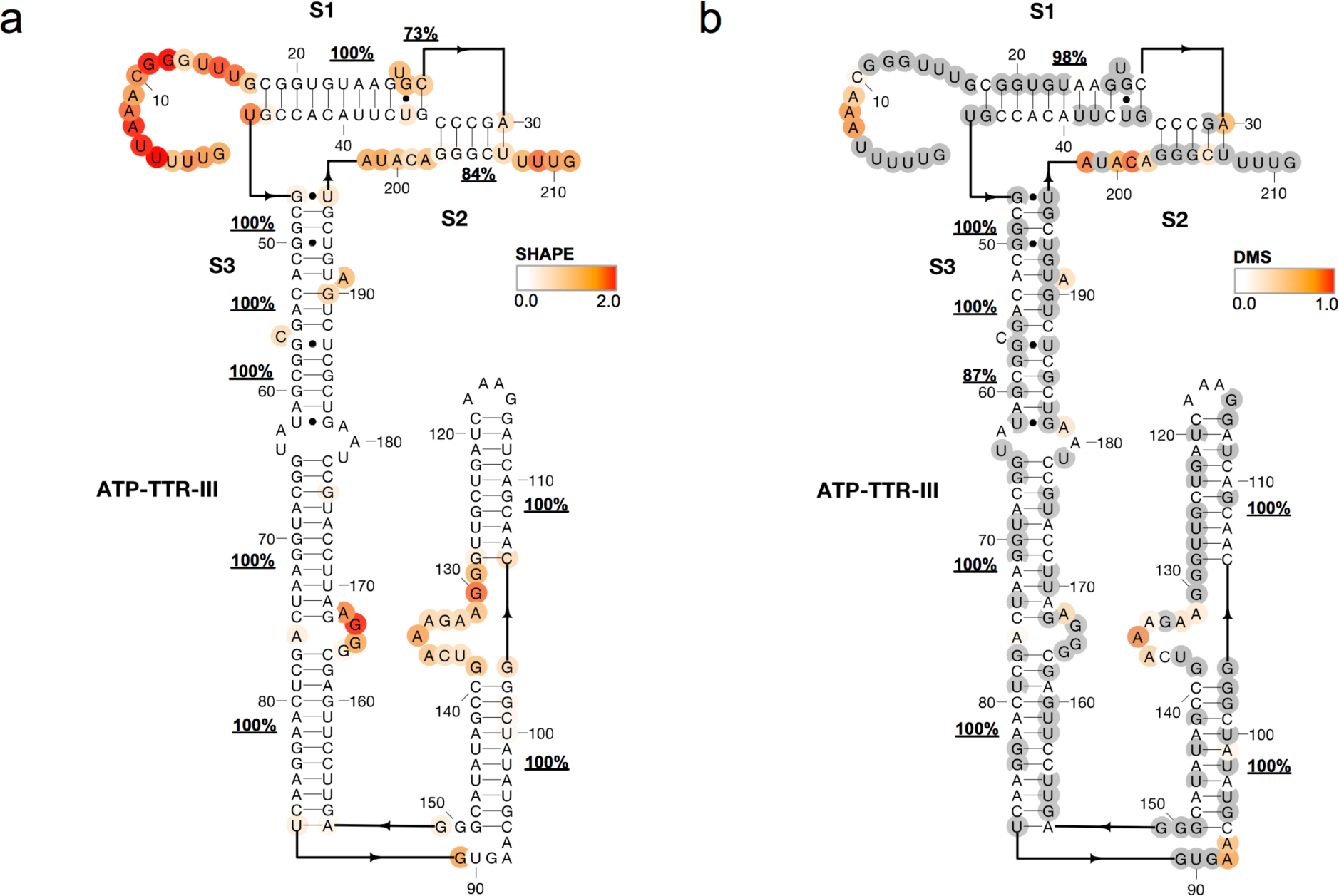
The secondary structure for FSE-ATP-TTR3, the SARS-CoV-2 frameshift stimulation element tagged by the ATP-TTR nanostructure as determined by 1D chemical SHAPE mapping, as computed in RNAStructure allowing for pseudoknots. Bootstrapping (100 iterations) support for each continuous helix is shown as an underlined percentage. Nucleotides are colored by (**a)** SHAPE reactivity or (**b**) dimethyl sulfate (DMS) reactivity. Bootstrapping probabilities are shown for the DMS case for helices that are also predicted with RNAstructure guided by DMS reactivity. Nucleotides unreactive to DMS are depicted in grey.

**Extended Data Fig. 11.**
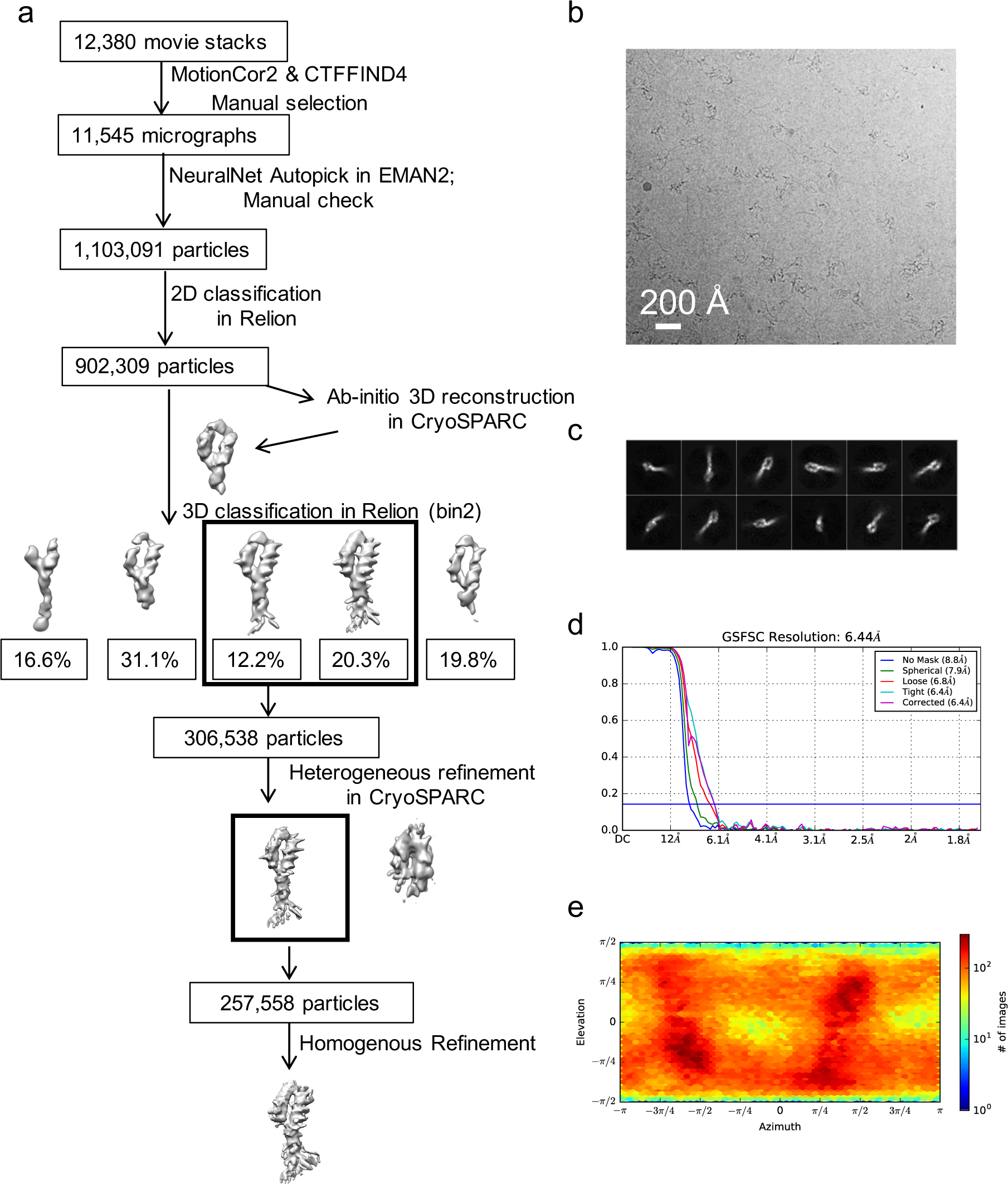
Single-particle cryo-EM analysis of the FSE-ATP-TTR3. **a.** Workflow of cryo-EM data processing of FSE. **b.** Representative motion-corrected cryo-EM micrograph with particles picked by the NeuralNet autopicking option in EMAN2. **c.** Reference-free 2D class averages. **d.** Gold standard FSC plots calculated in cryoSPARC. **e.** Euler angle distribution of the particle images.

**Extended Data Fig. 12.**
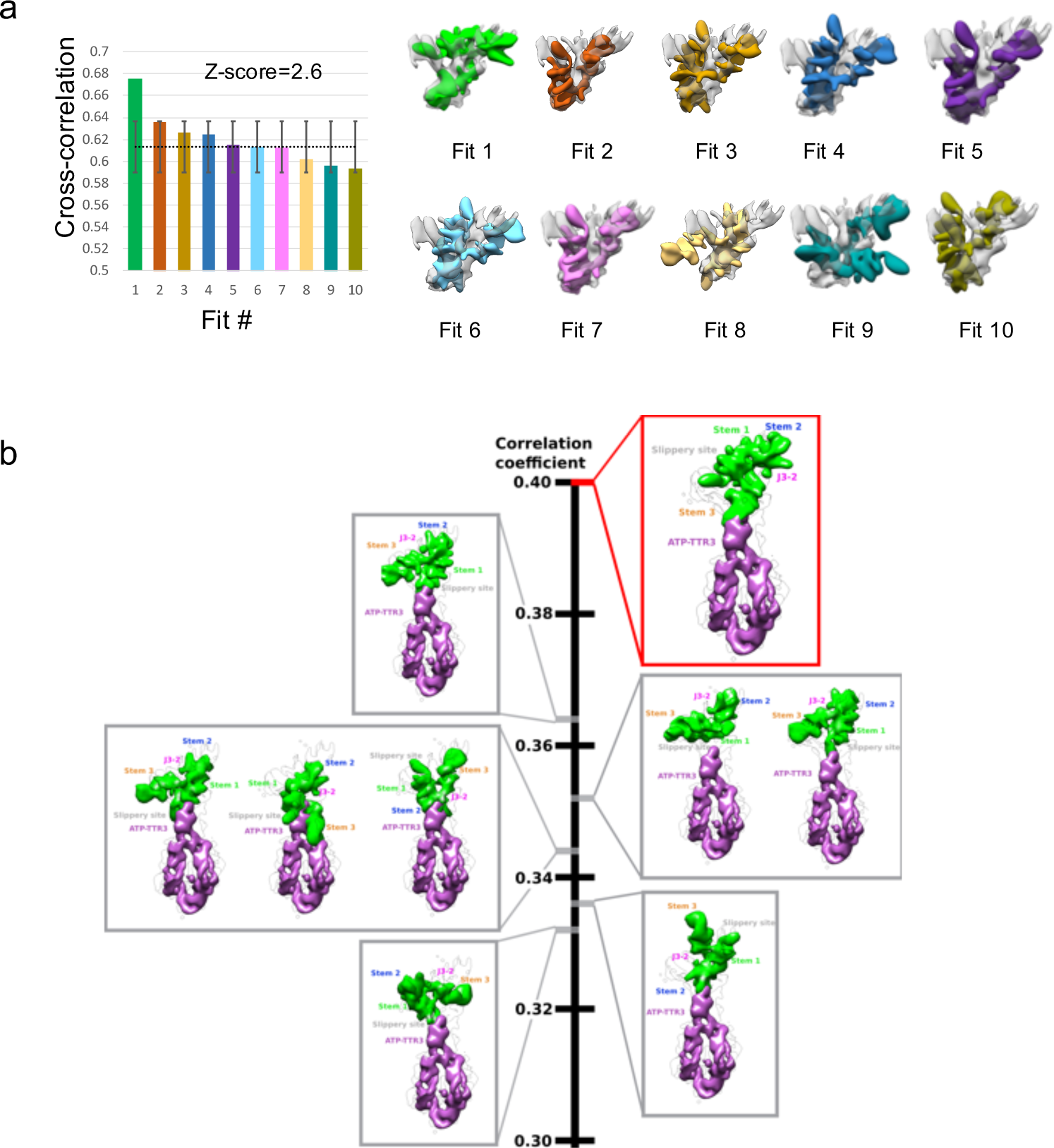
FSE-alone map fitting into the FSE-ATP-TTR3 map. The orientation of the FSE map was determined by conducting an exhaustive search of the FSE-alone map into the FSE-ATP-TTR3 map (with the ATP-TTR 3 part removed). The Fit to Segments option in the Segger plugin of Chimera was used. **a.** Fits with the top-ten cross-correlation (CC) scores are shown. The plot shows the top fit is significantly above the average of the other fits (Z-score = 2.6), indicating good confidence in the top fit as a correct placement. All 10 top fits are shown on the right. **b.** The exhaustive search was done using Situs, fitting the model of the FSE into the difference map between the FSE-ATP-TTR3 map and the ATP-TTR 3 map since Situs can only take a model as input to be fitted. The FSE-only map was then fitted into the same position as the model, and the fits of this map are visualized in red/gray boxes. The red box is for the top fit with the highest CC score, and this fit is identical to the top fit identified as shown in (a). The other gray boxes show fits of the model that have lower CC scores, which are plotted on the central bar.

**Extended Data Fig. 13.**
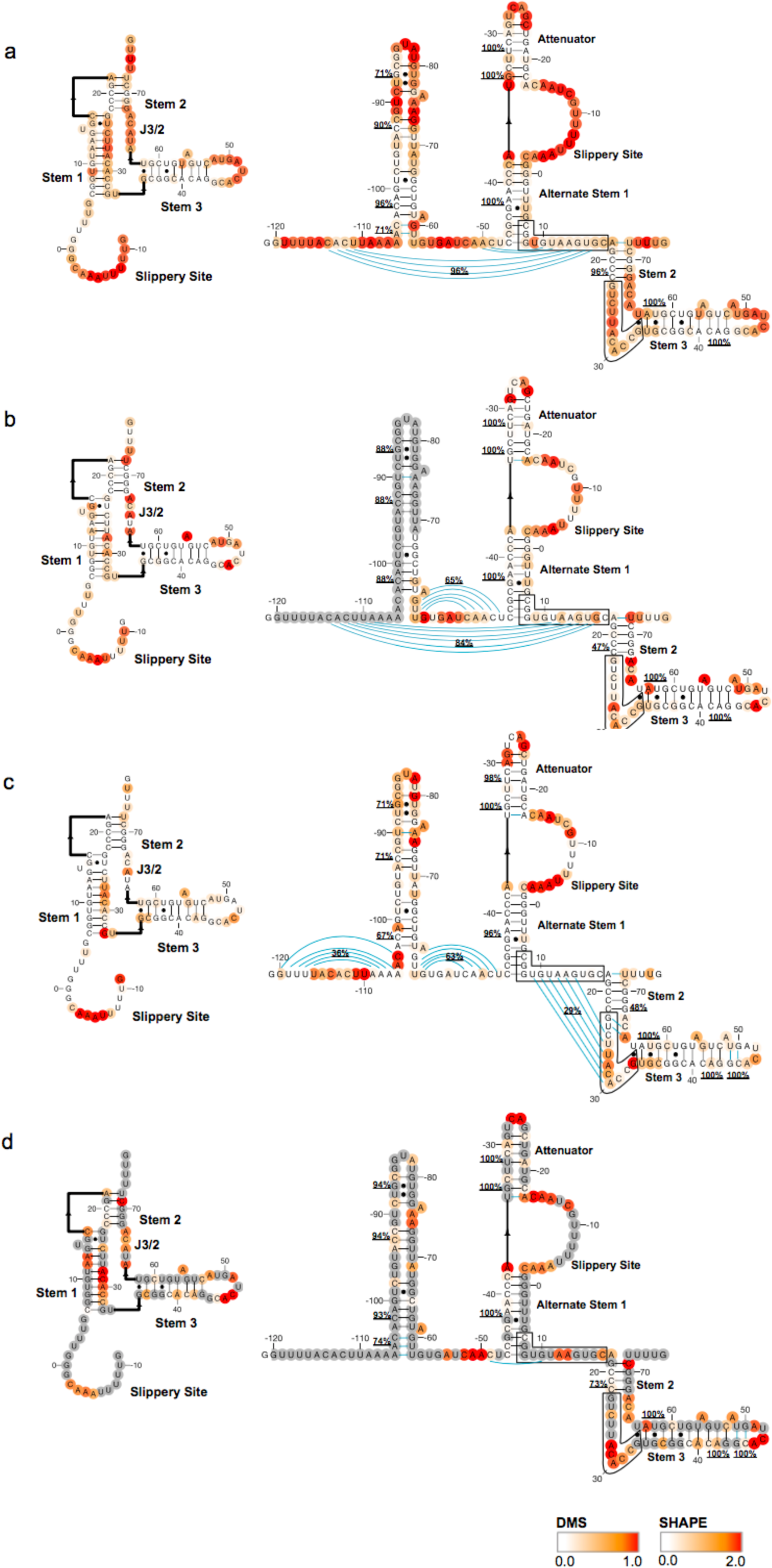
Secondary structure for the frameshift stimulation element (FSE) including 110 upstream nucleotides. Reactivity data are **a.** SHAPE reactivities collected in this work, **b.** SHAPE reactivities from Iserman, *et al*.^73^, **c.** SHAPE reactivities from Manfredonia, *et al*.^28^, and **d.** DMS reactivities from Lan *et al*^27^. We include reactivities superimposed on the FSE structure from Fig. 2 (left column) and reactivities superimposed on structures predicted using RNAstructure (right column). RNAstructure predictions are made with ShapeKnots if pseudoknots are predicted, and Fold otherwise. Bootstrapping (100 iterations) support for each helix is shown as an underlined percentage. Base pairs are shown in blue if they are not present in all four models. Boxed regions indicate strands of Stem 1 in the FSE structure.

**Extended Data Fig. 14.**
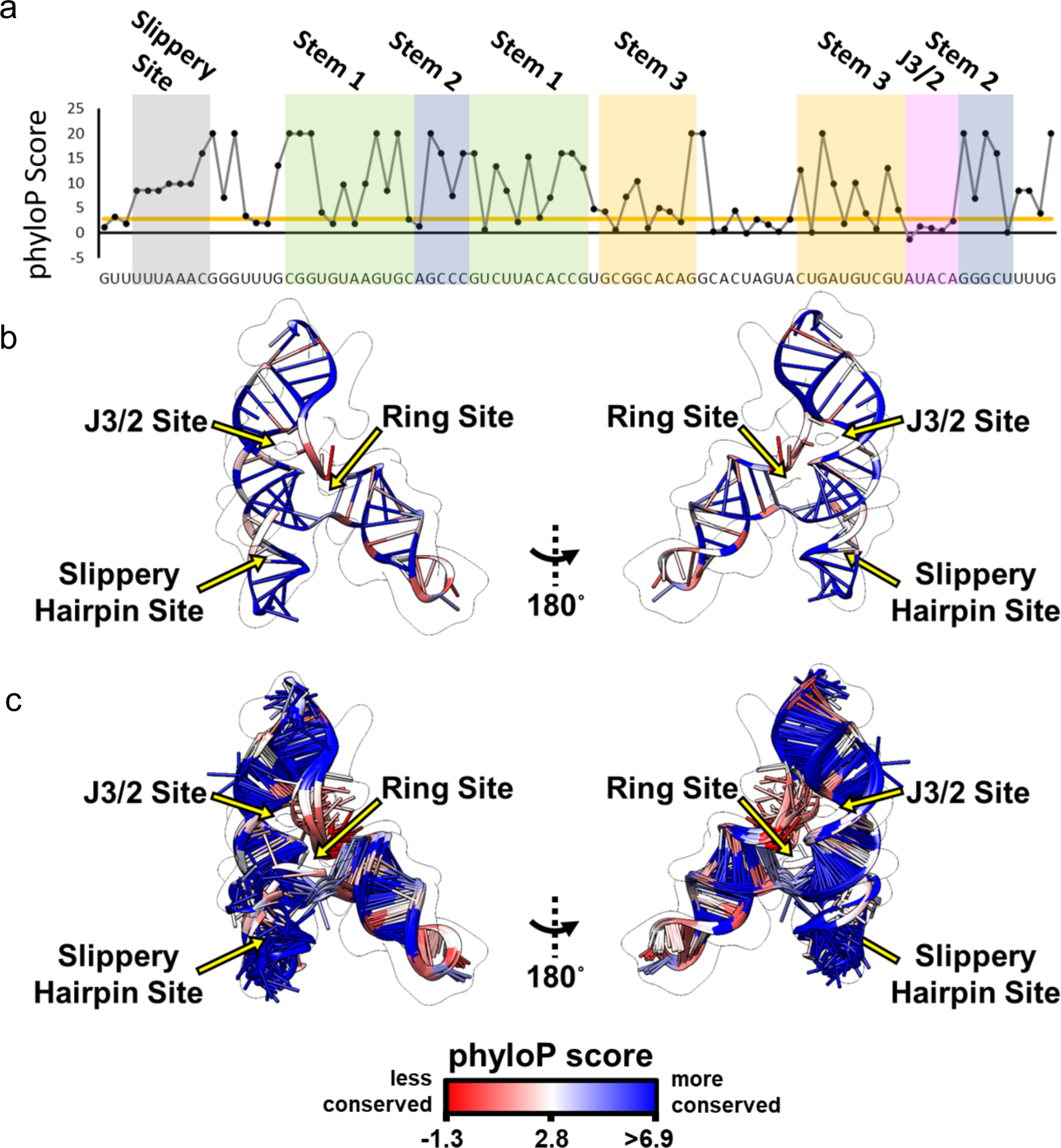
Evolutionary conservation of the SARS-CoV-2 FSE as determined by phyloP over 119 diverse coronavirus sequences^22^. **a.** Per-nucleotide phyloP scores (larger positive values indicate higher conservation; negative values indicate accelerated evolution^77^) of the FSE. The average phyloP score over the whole SARS-CoV-2 reference genome is shown as an orange line. **b-c.** the same model view shown in main text Fig. 4c but in ‘ribbon’ style (b) and with all ten models from the autoDRRAFTER modelling (c). Structures are colored by phyloP scores so that blue vs. red highlight regions of higher vs. lower conservation than the average genomic phyloP score, respectively.

## Movie legends

**Extended Data Movie 1**. Cryo-EM map-derived atomic model of 28-kDa Frameshift stimulation element (FSE) from the SARS-CoV-2 RNA Genome.

**Extended Data Movie 2**. FSE-alone map fitting into the FSE-ATP-TTR3 map. The orientation of the FSE map was determined by conducting an exhaustive search of the FSE-alone map into the FSE-ATP-TTR3 map.

