## Extended Data Table 1 for "Cryo-electron Microscopy and Exploratory Antisense Targeting of the 28-kDa Frameshift Stimulation Element from the SARS-CoV-2 RNA Genome"

### Cryo-EM data collection and validation statistics

| | FSE (EMDB-22296) (PDB 6XRZ) | FSE-ATP-TTR3 (EMDB-22297) |
|---|---|---|
| **Data collection and processing** | | |
| Magnification | 165k | 165k |
| Voltage (kV) | 300 | 300 |
| Electron exposure (e–/Å$^2$) | 8.3 | 8.3 |
| Defocus range (μm) | -1.2 - -3.5 | -1.2 - -3.5 |
| Pixel size (Å) | 0.82 | 0.82 |
| Symmetry imposed | C1 | C1 |
| Initial particle images (no.) | 1,063,711 | 1,103,091 |
| Final particle images (no.) | 109,137 | 257,558 |
| Map resolution (Å) | 6.9 | 6.4 |
| FSC threshold | 0.143 | 0.143 |
| **Validation** | | |
| MolProbity score | 2.87 | N/A |
| Clashscore | 16.73 | N/A |
| Poor rotamers (%) | N/A | N/A |
| RNA Geometry | | |
| Bad bonds | 0 / 2085 | N/A |
| Bad angles | 5 / 3246 | N/A |
| Probably wrong sugar puckers | 4 | N/A |
