## Extended Data Table 2 for "Cryo-electron Microscopy and Exploratory Antisense Targeting of the 28-kDa Frameshift Stimulation Element from the SARS-CoV-2 RNA Genome"

| Full sequences | |
|---|---|
| **Name** | **Sequence** |
| **SARS-CoV-2_FSE_13459-13546** (+Phi2.5 T7 RNAP promoter, underlined) | **DNA:** TTCTAATACGACTCACTATT<br>GTTTTTAAACGGGTTTGCGGTGTAAGTGCAGCCCGTCTTACACCGTG<br>CGGCACAGGCACTAGTACTGATGTCGTATACAGGGCTTTTG<br>**RNA:**<br>GUUUUUAAACGGGUUUGCGGUGUAAGUGCAGCCCGUCUUACAC<br>CGUGCGGCACAGGCACUAGUACUGAUGUCGUAUACAGGGCUUU<br>UG |
| **SARS-CoV-2_FSE_1D-chemical-mapping** (+Phi6.5 T7 RNAP promoter, underlined; reference hairpins, italics; Tail2 RT binding site, underlined italics) | **DNA:**<br>TTCTAATACGACTCACTATA *GGTCAGCGAGTAGCTGAC*<br>GTTTTTAAACGGGTTTGCGGTGTAAGTGCAGCCCGTCTTACACCGTG<br>CGGCACAGGCACTAGTACTGATGTCGTATACAGGGCTTTTG<br>*GTCAGCGAGTAGCTGAC* *AAAGAAACAACAACAACAAC*<br>**RNA:**<br>*GGUCAGCGAGUAGCUGAC*<br>GUUUUUAAACGGGUUUGCGGUGUAAGUGCAGCCCGUCUUACAC<br>CGUGCGGCACAGGCACUAGUACUGAUGUCGUAUACAGGGCUUU<br>UG *GUCAGCGAGUAGCUGAC* *AAAGAAACAACAACAACAAC* |
| **SARS-CoV-2_plusFSE_13349-13546** (+Phi6.5 T7 RNAP promoter, underlined) | **DNA:** TTCTAATACGACTCACTATA<br>GGTTTTACACTTAAAAACACAGTCTGTACCGTCTGCGGTATGTGGAA<br>AGGTTATGGCTGTAGTTGTGATCAACTCCGCGAACCCATGCTTCAGT<br>CAGCTGATGCACAATCGTTTTTAAACGGGTTTGCGGTGTAAGTGCAG<br>CCCGTCTTACACCGTGCGGCACAGGCACTAGTACTGATGTCGTATAC<br>AGGGCTTTTG<br>**RNA:**<br>GGUUUUACACUUAAAAACACAGUCUGUACCGUCUGCGGUAUGU<br>GGAAAGGUUAUGGCUGUAGUUGUGAUCAACUCCGCGAACCCAU<br>GCUUCAGUCAGCUGAUGCACAAUCGUUUUUAAACGGGUUUGCG<br>GUGUAAGUGCAGCCCGUCUUACACCGUGCGGCACAGGCACUAGU<br>ACUGAUGUCGUAUACAGGGCUUUUG |
| **SARS-CoV-2_plusFSE_1D-chemical-mapping** (+Phi6.5 T7 RNAP promoter, underlined; reference hairpins, italics; Tail2 RT binding site, underlined italics) | **DNA:**<br>TTCTAATACGACTCACTATA *GGTCAGCGAGTAGCTGAC*<br>GGTTTTACACTTAAAAACACAGTCTGTACCGTCTGCGGTATGTGGAA<br>AGGTTATGGCTGTAGTTGTGATCAACTCCGCGAACCCATGCTTCAGT<br>CAGCTGATGCACAATCGTTTTTAAACGGGTTTGCGGTGTAAGTGCAG<br>CCCGTCTTACACCGTGCGGCACAGGCACTAGTACTGATGTCGTATAC<br>AGGGCTTTTG *GUCAGCGAGUAGCUGAC*<br>*AAAGAAACAACAACAACAAC*<br>**RNA:**<br>*GGUCAGCGAGUAGCUGAC*<br>GGUUUUACACUUAAAAACACAGUCUGUACCGUCUGCGGUAUGU<br>GGAAAGGUUAUGGCUGUAGUUGUGAUCAACUCCGCGAACCCAU<br>GCUUCAGUCAGCUGAUGCACAAUCGUUUUUAAACGGGUUUGCG<br>GUGUAAGUGCAGCCCGUCUUACACCGUGCGGCACAGGCACUAGU<br>ACUGAUGUCGUAUACAGGGCUUUUG *GUCAGCGAGUAGCUGAC*<br>*AAAGAAACAACAACAACAAC* |
| **FSE-dl-ATP-TTR-3** (+Phi2.5 T7 RNAP promoter, underlined; FSE black, ATP-TTR-3, grey) | **DNA:** TTCTAATACGACTCACTATT<br>GTTTTTAAACGGGTTTGCGGTGTAAGTGCAGCCCGTCTTACACCGTG<br>CGGCACAG<br>CGGCGATATGGCATGGAATCAGCTCAAGGAACTGTGAACGTATATC |

| | |
|---|---|
| | GGGCAACGACTAGGAAACTAGTCGTTGGGAAGAAACTGCCGATATA<br>CGGGAGTTCCTTGAGCGGGAGATTCCATGCCTAAGTCGC<br>TCTGATGTCGTATACAGGGCTTTTG<br>**RNA:**<br>GUUUUUAAACGGGUUUGCGGUGUAAGUGCAGCCCGUCUUACAC<br>CGUGCGGCACAG<br>CGGCGAUAUGGCAUGGAAUCAGCUCAAGGAACUGUGAACGUAUA<br>UCGGGCAACGACUAGGAAACUAGUCGUUGGGAAGAAACUGCCGA<br>UAUACGGGAGUUCCUUGAGCGGGAGAUUCCAUGCCUAAGUCGC<br>UCUGAUGUCGUAUACAGGGCUUUUG |
| **FSE-dl-ATP-TTR-3_1D-chemical-mapping** (+Phi6.5 T7 RNAP promoter, underlined; reference hairpins, italics; Tail2 RT binding site, underlined italics; FSE black, ATP-TTR-3, grey) | **DNA:** TTCTAATACGACTCACTATA *GGTCAGCGAGTAGCTGACAG*<br>GTTTTTAAACGGGTTTGCGGTGTAAGTGCAGCCCGTCTTACACCGTG<br>CGGCACAG<br>CGGCGATATGGCATGGAATCAGCTCAAGGAACTGTGAACGTATATC<br>GGGCAACGACTAGGAAACTAGTCGTTGGGAAGAAACTGCCGATATA<br>CGGGAGTTCCTTGAGCGGGAGATTCCATGCCTAAGTCGC<br>TCTGATGTCGTATACAGGGCTTTTG *AGTCAGCGAGTAGCTGAC*<br>*AAAGAAACAACAACAACAAC*<br>**RNA:** *GGUCAGCGAGUAGCUGACAG*<br>GUUUUUAAACGGGUUUGCGGUGUAAGUGCAGCCCGUCUUACAC<br>CGUGCGGCACAG<br>CGGCGAUAUGGCAUGGAAUCAGCUCAAGGAACUGUGAACGUAUA<br>UCGGGCAACGACUAGGAAACUAGUCGUUGGGAAGAAACUGCCGA<br>UAUACGGGAGUUCCUUGAGCGGGAGAUUCCAUGCCUAAGUCGC<br>UCUGAUGUCGUAUACAGGGCUUUUG *AGUCAGCGAGUAGCUGAC*<br>*AAAGAAACAACAACAACAAC* |

| PCR assembly primers | |
|---|---|
| **SARS-CoV-2_FSE_Phi2.5** | |
| **FSE_p2-5_1F** | TTCTAATACGACTCACTATTGTTTTTAAACGGGTTTGC |
| **FSE_p2-5_2R** | AGACGGGCTGCACTTACACCGCAAACCCGTTTAAAAACAATAGTGA<br>G |
| **FSE_p2-5_3F** | AGTGCAGCCCGTCTTACACCGTGCGGCACAGGCACTAGTACTGAT |
| **FSE_p2-5_4R** | CAAAAGCCCTGTATACGACATCAGTACTAGTGCCTGTGCCGCACG |
| **SARS-CoV-2_FSE_1D-chemical-mapping_Phi6.5** | |
| **FSE_T7_CM-1F** | TTCTAATACGACTCACTATAGGTCAGCGAGTAGCTGACGTTTTTAAA<br>CGGGTTTGCGGTGTAAGTGCAGCCCGTCTTACACCGTGCGGCACA |
| **FSE_T7_CM-2R** | GTTGTTGTTGTTGTTTCTTTGTCAGCTACTCGCTGACCAAAAGCCCTG<br>TATACGACATCAGTACTAGTGCCTGTGCCGCACGGTGTAAGAC |
| **SARS-CoV-2_plusFSE_Phi6.5** | |
| **plusFSE_p6.5-1F** | TTCTAATACGACTCACTATAGGTTTTACACTTAAAAACA |
| **plusFSE_p6.5-2R** | ACATACCGCAGACGGTACAGACTGTGTTTTTAAGTGTAAAACCTATA<br>GTGAGTCGTATT |
| **plusFSE_p6.5-3F** | ACCGTCTGCGGTATGTGGAAAGGTTATGGCTGTAGTTGTGATCAACT<br>CCGCGAACCCA |
| **plusFSE_p6.5-4R** | GCAAACCCGTTTAAAAACGATTGTGCATCAGCTGACTGAAGCATGG<br>GTTCGCGGAGTTGA |
| **plusFSE_p6.5-5F** | TCGTTTTTAAACGGGTTTGCGGTGTAAGTGCAGCCCGTCTTACACCG<br>TGCGGC |

| | |
|---|---|
| **plusFSE_p6.5-6R** | AAAAGCCCTGTATACGACATCAGTACTAGTGCCTGTGCCGCACGGTGTAAGA |
| **SARS-CoV-2_plusFSE_1D-chemical-mapping_Phi6.5** | |
| **plusFSE_T7_CM-1F** | TTCTAATACGACTCACTATAGGTCAGCGAGTAGCTGACGGTTTTACACTTAAAAACACAGTCTGTACCGTCTGCGGTATGTGGAAAGGTTATGGC |
| **plusFSE_T7_CM-2R** | CGCAAACCCGTTTAAAAACGATTGTGCATCAGCTGACTGAAGCATGGGTTCGCGGAGTTGATCACAACTACAGCCATAACCTTTCC |
| **plusFSE_T7_CM-3F** | CGGGTTTGCGGTGTAAGTGCAGCCCGTCTTACACCGTGCGGCACAGGCACTAGTACTGATGTCGTATACAGGGCTTTTGGT |
| **plusFSE_T7_CM-4R** | GTTGTTGTTGTTGTTTCTTTGTCAGCTACTCGCTGACCAAAAGCCCT |
| **FSE-dl-ATP-TTR-3_Phi2.5** | |
| **FSE-dl-ATP-TTR-3_p2-5-1F** | TTCTAATACGACTCACTATTGTTTTTAAACGGGTTTGCGGTGTAAGTGCAGCCCGTCTTACACCGTGCGGCACAGCGGCG |
| **FSE-dl-ATP-TTR-3_p2-5-2R** | CCTAGTCGTTGCCCGATATACGTTCACAGTTCCTTGAGCTGATTCCATGCCATATCGCCGCTGTGCCGCACG |
| **FSE-dl-ATP-TTR-3_p2-5-3F** | CGGGCAACGACTAGGAAACTAGTCGTTGGGAAGAAACTGCCGATATACGGGAGTTCCTTGAGCGGGAGATTCCATGCCTAAGTCGCTCT |
| **FSE-dl-ATP-TTR-3_p2-5-4R** | CAAAAGCCCTGTATACGACATCAGAGCGACTTAGGCATGGAATCT |
| **FSE-dl-ATP-TTR-3_1D-chemical-mapping_Phi6.5** | |
| **FSE-dl-ATP-TTR-3_T7_CM-1F** | TTCTAATACGACTCACTATAGGTCAGCGAGTAGCTGACAGGTTTTTAAACGGGTTTGCGGTGTAAGTGCAGCCCGTCTTACACCGTGCGGCACAGCGGCG |
| **FSE-dl-ATP-TTR-3_T7_CM-2R** | CCTAGTCGTTGCCCGATATACGTTCACAGTTCCTTGAGCTGATTCCATGCCATATCGCCGCTGTGCCGCACG |
| **FSE-dl-ATP-TTR-3_T7_CM-3F** | CGGGCAACGACTAGGAAACTAGTCGTTGGGAAGAAACTGCCGATATACGGGAGTTCCTTGAGCGGGAGATTCCATGCCTAAGTCGCTCT |
| **FSE-dl-ATP-TTR-3_T7_CM-4R** | GTTGTTGTTGTTGTTTCTTTGTCAGCTACTCGCTGACTCAAAAGCCCTGTATACGACATCAGAGCGACTTAGGCATGGAATCT |

| **2D Chemical Mapping Primers (M2-seq)** | |
|---|---|
| **Error Prone PCR Primers** | |
| **Forward** | TTCTAATACGACTCACTATAGGUCAGC |
| **Reverse** | GTTGTTGTTGTTGTTTCTTTGTCAGC |
| **Retrotranscription Primer** | |
| **RTB retrotranscription primer** (*s indicate index nucleotides, annealing nucleotides bolded) | AATGATACGGCGACCACCGAGATCTACACTCTTTCCCTACACGACGCTCTTCCGATCT ************ **GTTGTTGTTGTTGTTTCTTT** |
| **Second Strand Synthesis Primer** | |
| **Second strand synthesis primer** (annealing nucleotides bolded) | CAAGCAGAAGACGGCATACGAGATCGGTCTCGGCATTCCTGCTGAACCGCTCTTCCGATCT **GGTCAGC** |
| **iTru Primers** | |
| **p5** (* indicates phosphorothioate bond) | AATGATACGGCGACCACCGAGA*T |
| **p7** (* indicates phosphorothioate bond) | CAAGCAGAAGACGGCATACGAGA*T |
| **Retrotranscription Primer Indices** | |
| **RTB-001** | GAGGCCTTGGCC |
| **RTB-002** | CTTTAAAATATA |
| **RTB-003** | TGACTTGCACAT |

| | |
|---|---|
| **RTB-004** | TGCGCCATTGCT |
| **RTB-005** | ACAAAATGGTGG |
| **RTB-006** | CTGCGTGCAAAC |

| **Full secondary structures** | |
|---|---|
| **Name** | **Secondary structure** |
| **SARS-CoV-2_FSE_13459-13546** | **Literature:**<br>..............((((((((((...[[[[[[[))))))))))))((((((((.........))).))))))...]].]]]]]....<br>**1D chemical mapping, 1M7 with pseudoknots:**<br>.((((((.....))))))................(((((((((.(([[[[[[)))))))))))).(((((((((.........))).))))))<br>.....]]]]]....((((((.....))))))<br>**1D chemical mapping, 1M7 without pseudoknots:**<br>.((((((.....)))))).........(.((((.((((((((((.((.....))))))))))))))((((((((((((....)).))).)))))<br>)......))))...)((((((.....))))))<br>**1D chemical mapping, DMS with pseudoknots:**<br>.((((((.....))))))..........(((((...[[[[[[[[[....))))).]]]]]]]]].((((((((((((....)).))).)))))<br>)................((((.....))))..<br>**1D chemical mapping, DMS without pseudoknots:**<br>.((((((.....))))))..........(((((.((((((((((.((.....))))))))))))))((((((((((((....)).))).)))))<br>)......)))))......((((.....))))..<br>**2D chemical mapping, + 1D DMS with pseudoknots**<br>...............((((((((((....[[[[..))))))))))((((((((.........))).)))))).....]]]].....|
| **SARS-CoV-2_plusFSE_13349-13546** | **Literature:**<br><br>**1D chemical mapping, 1M7 with pseudoknots:**<br>.......((((.....((((((((...(((..((((.....))))...)))...)))))).))........((.((((((((((..((.((((<br>....)))).))................))))))))).)).)))).(((((..........((((((((((.........))).)))))))).<br>..)))))....<br>**1D chemical mapping, 1M7 without pseudoknots:**<br>.((((((.....))))))........((((.....((((((((...(((..(.((.....)).)...)))...)))))).)).(....).((.((<br>((((((((..((.((((....)))).)).(....).........)))))))))).)).)))).(((((..........(((((((((((....<br>.....))).))))))))...)))))....((((((.....))))))<br>**1D chemical mapping, DMS with pseudoknots:**<br>..((((((.....))))).(((.....)))..........((((...((((..((((.....(((........(((((((.(((.........(((<br>(((((((.(((.((((....)))).)))...............))))))))))..))).))))))))..)))....))))))))...))))...<br>....((((((((.........)))......)))))............<br>**1D chemical mapping, DMS without pseudoknots:**<br>..((((((.....))))).(((.....)))..........((((.(.((((..((((.....(((........(((((((.(((........(.(((<br>(((((((.(((.((((....)))).)))...............))))))))))).)))).))))))))..)))....))))))))).).))))..<br>.....((((.(((.........)))..)))).((((.....))))..<br>**2D chemical mapping, + 1D DMS with pseudoknots**<br>..............(((((((((...(((.((((((...))))))..)))...))))))).)))..........((((((((((.(((.((((<br>....)))).)))...............))))))))))..........(((((..........(((((((((((((....)).))).)))))))))).<br>..)))))....<br>**1D chemical mapping, 1M7 with pseudoknots, Manfredonia et al. 2020 bioRxiv**<br>.((((((......)))).)((((((...(((.(((((.....)))).).)))...))))))(((((....)))))((((((((((.(((.((<br>((....)))).)))...............))))))))))[[[[[[...(((((((]]]]]]..(((((((((((((....)).))).))))))<br>))))).))))))....|

| | |
|---|---|
| | **1D chemical mapping, 1M7 without pseudoknots, Iserman et al. 2020 bioRxiv**<br>.......((((((.....((((((...(((.(((((.....)))))..)))...))))))(((((....)))))((((((((((.(((.(((<br>(....)))).)))...............))))))))))..)))))).(((((..........((((((((((.........))).))))))))<br>...)))))....<br>**1D chemical mapping, DMS without pseudoknots, Lan et al. 2020 bioRxiv**<br>.......(((.....(((((((((...(((.(((((.....)))))..)))...)))))).)))))).....(.((((((((((.(((.(((<br>(....)))).)))...............))))))))))..)........((((..........((((((((((((....)).))).))))))))<br>...)))).....<br> |
| **FSE-dl-ATP-TTR-3** | **Literature:**<br>...............((((((((((...[[[[[[[))))))))))))(((((((((.((((((..((((((((((((.((((((((((((...<br>..((((((((((.((((((((((....))))))))))...........))))))))))..))))))))))))....)))))))))))...)<br>))))))))).)))))).]].]]]]]....<br>**1D chemical mapping, 1M7 with pseudoknots:**<br>.((((((.....))))))...................(((((((((.(([[[[[[))))))))))).(((((((((.((((((..(((((((<br>((((.((((((((((((.....((((((((((.((((((((((....))))))))))...........))))))))))..))))))))))<br>))....))))))))))))...))))))))).)))))).....]]]]].....((((((.....))))))<br>**1D chemical mapping, 1M7 without pseudoknots:**<br>.((((((.....))))))...................(((((((((.((.....))))))))))).(((((((((.((((((..((((((((<br>(((.((((((((((((.....((((((((((.((((((((((....))))))))))(.......)..))))))))))..)))))))))))<br>)....))))))))))))...))))))))).)))))).................((((((.....))))))<br>**1D chemical mapping, DMS with pseudoknots:**<br>...((((.....((((((((((((...((((((...[[[[[[[[....))))))]]]]]]]]].(((((((((.((((((..((((((<br>(((((.((((((((((((.....((((((((((.((((((((((....))))))))))...((...)).))))))))))..)))))))))<br>)))....))))))))))))...))))))))).)))))).....)))))))....)))))).....))))..<br>**1D chemical mapping, DMS without pseudoknots:**<br>.((((((.....))))))..((......)).((((.((((((((((.((.....))))))))))))))(((((((((.((((((..(((((((<br>((((.((((((((((((.....((((((((((.((((((((((....))))))))))...((...)).))))))))))..))))))))))<br>))....))))))))))))...))))))))).))))))......))))......(((((.....))))).<br>**2D chemical mapping, + 1D DMS with pseudoknots**<br>.........[[[[[[..((((((((((....]]]]]]))))))))))(((((((((.((((((..((((((((((((.((((((((((((.<br>....((((((((((.((((((((((....))))))))))...........))))))))))..))))))))))))....))))))))))))..<br>.))))))))).)))))).............. |
